# Connectomic identification and three-dimensional color tuning of S-OFF midget ganglion cells in the primate retina

**DOI:** 10.1101/482653

**Authors:** Lauren E Wool, Orin S Packer, Qasim Zaidi, Dennis M Dacey

## Abstract

In the trichromatic primate retina, the ‘midget’ retinal ganglion cell is the classical substrate for red-green color signaling, with a circuitry that enables antagonistic responses between long (L)- and medium (M)-wavelength sensitive cone inputs. Previous physiological studies show that some OFF midget ganglion cells may receive sparse input from short (S)-wavelength sensitive cones, but the effect of S-cone inputs on the chromatic tuning properties of such cells has been unexplored. Moreover, anatomical evidence for a synaptic pathway from S cones to OFF midget ganglion cells through OFF-midget bipolar cells remains ambiguous. In this study we address both questions for the macaque monkey retina. First, we used serial block-face electron microscopy (SBEM) to show that every S-cone in the parafoveal retina synapses principally with a single OFF-midget bipolar cell which in turn forms a private-line connection with an OFF midget ganglion cell. Second, we used patch electrophysiology to characterize the chromatic tuning of OFF midget ganglion cells in the near peripheral retina that receive combined input from L, M and S cones. These ‘S-OFF’ midget cells have a characteristic S-cone spatial signature, but demonstrate heterogeneous color properties due to variable strength of L, M, and S cone input across the receptive field. Together these findings strongly support the hypothesis that the OFF midget pathway is the major conduit for S-OFF signals in primate retina, and redefines the pathway as a chromatically complex substrate that encodes color signals beyond the classically recognized L vs. M and S vs. L+M cardinal mechanisms.

**Significance statement:** The first step of color processing in the visual pathway of primates occurs when signals from short- (S), middle- (M) and long- (L) wavelength sensitive cone types interact antagonistically within the retinal circuitry to create color-opponent pathways. The midget (L vs. M or ‘red-green’) and small bistratified (S vs. L+M, or ‘blue-yellow’) appear to provide the physiological origin of the cardinal axes of human color vision. Here we confirm the presence of an additional S-OFF midget circuit in the macaque monkey fovea with scanning block-face electron microscopy (SBEM) and show physiologically that a subpopulation of S-OFF midget cells combine S, L and M cone inputs along non-cardinal directions of color space, expanding the retinal role in color coding.

## Introduction

In humans and most non-human primates, color processing begins in the retinal circuitry where signals arising from three spectrally distinct photoreceptor types (the long (L), middle (M), and short (S) wavelength sensitive cones), interact antagonistically. Two anatomically distinct circuits correlate strongly with cardinal directions in color space classically identified in human psychophysics (Krauskopf et al., 1982): a ‘red-green’ mechanism, which combines antagonistic inputs from L and M cones, and served by the midget ganglion cell (Dacey and Lee, 1994; Crook et al., 2011) and a ‘blue-yellow’ mechanism in which S cones oppose a combined L+M cone signal and served by the S-ON small bistratified ganglion cell (Dacey and Lee, 1994). The midget L vs M pathway includes ON and OFF cell populations responding positively to increments and decrements respectively. The S-ON pathway has been characterized in detail (Mariani, 1984; Kouyama and Marshak, 1992; Calkins et al., 1998; Herr et al., 2003; Crook et al., 2009) but a complementary S-OFF circuit for transmission of Scone decrements has remained elusive.

Two pieces of evidence point to an anatomically and physiologically distinct subpopulation of OFF type midget ganglion cells as the origin of the S-OFF chromatic pathway (Dacey et al., 2014). First, elegant serial section electron microscopy from the macaque monkey fovea showed that OFF midget bipolar cells made basal, OFF-type contacts with presumed S-cones and would thus clearly comprise an S-OFF midget pathway (Klug et al., 2003). Second, midget ganglion cells in the retinal periphery, especially the OFF-type, can combine inputs of the same sign from S, L and M cone types (Field et al., 2010).

In the greater scope of S-cone circuitry, the significance of these two results remains unclear. S-OFF cells (Valberg et al., 1986) recorded at the level of the lateral geniculate nucleus show spatiotemporal properties that argue strongly against their basis in the midget pathway (Tailby et al., 2008a; Tailby et al., 2008b). More fundamentally, how—or even whether—additional weak S input to peripheral midget cells effects their color tuning properties is unknown. Moreover, the striking absence of an S-OFF midget pathway has been demonstrated in a marmoset (Lee et al., 2005), despite the clear identification of a homologous S-ON pathway circuitry in this New World monkey (Ghosh et al., 1997). The lack of an S-OFF midget pathway in marmoset echoes an earlier serial section electron microscopic study in human retina that also failed to observe the presence of an OFF midget bipolar contact at the presumed S cone pedicle (Kolb et al., 1997). Correspondingly, the lack of an ERG d-wave S-OFF signature has been taken as evidence for the absence of an S-OFF pathway in the human retina (Maguire et al., 2018). Finally, it has been shown that a sign-inverting amacrine cell circuit (Chen and Li, 2012) can be utilized to create an S-OFF pathway in a ground squirrel (Sher and DeVries, 2012). Thus, both the anatomical origins and functional properties of an S-OFF color opponent pathway remain unsettled.

Here we apply serial block-face electron microscopy (SBEM) to the macaque monkey fovea to show that all S cones synapse with a single OFF midget bipolar cell, which in turn makes a private-line connection with an OFF midget ganglion cell, confirming and extending the earlier result (Klug et al., 2003). Additionally, through single-cell patch recordings, we report OFF midget ganglion cells in the near retinal periphery that receive S input and exhibit heterogeneous color properties due to combined inputs from L, M, and S cones. Our results advance the hypothesis that the OFF midget pathway is the principal conduit for S-OFF signals in macaque retina.

Moreover, this work identifies a chromatically complex substrate at the retinal level that can encode color signals distinct from the cardinal L vs. M and S vs. L+M axes.

## Materials and methods

### Retinal isolation

Eyes were removed from deeply anesthetized male and female macaque monkeys at the time of euthanasia (*Macaca nemestrina, Macaca fascicularis*, or *Macaca mulatta*) via the Tissue Distribution Program of the Washington National Primate Research Center and in accordance to protocols reviewed and approved by UW’s Institutional Animal Care and Use Committee (IACUC). After enucleation, the anterior chamber of the eye was removed, the vitreous drained, and the remaining eyecup placed in oxygenated Ames medium (A1420, Sigma). The choroid was then carefully dissected from the sclera to maintain intimate contact between neural retina, retinal pigment epithelium and choroid.

### Serial block-face electron microscopy

Tissue was first isolated as described above. After isolation, the central retina (including the fovea) was dissected and placed in a 4% glutaraldehyde solution for approximately two hours of fixation. The foveal tissue was then rinsed thoroughly in cacodylate buffer (0.1M, pH 7.4) and incubated in a 1.5% potassium ferrocyanide and 2% osmium tetroxide (OsO_4_) solution in 0.1M cacodylate buffer for one hour. After washing, the tissue was placed in a freshly made thiocarbohydrazide solution (0.1g TCH in 10 ml double-distilled H_2_O heated to 600°C for 1 h) for 20 min at room temperature (RT). After another rinse at RT, the tissue was incubated in 2% OsO_4_ for 30 min at RT. The samples were rinsed again and stained *en bloc* in 1% uranyl acetate overnight at 40°C, washed and stained with Walton’s lead aspartate for 30 min. After a final wash, the retinal pieces were dehydrated in a graded alcohol series, and placed in propylene oxide at RT for 10 min. The tissue was then embedded in Durcupan resin. Semi-thin vertical sections through the retinal layers (0.5–1 μm thick) were cut and stained with toluidine blue and examined to determine the location of the foveal center. A region of interest was chosen on the foveal slope ~400 microns from the foveal center for block-face imaging in the scanning electron microscope (SEM). The block was then trimmed, gold-coated by standard methods and mounted in a GATAN/ Zeiss (3View) SEM. The block face was imaged in an array of 25 40 μm × 40 μm tiles (10% overlap between tiles) that extended from the Henle fiber layer to the optic fiber layer (~200 microns vertical and lateral extent (see Figure 1A for an image of most of the sampled area). The block face was imaged after each of 420 sections cut at 80 nm thickness. Scanning was performed with a 5 nm X-Y resolution and a dwell time of 1 μsec. The resulting set of 10,500 tiff images were contrast-normalized, stitched into 420 layers, then aligned into a volume using procedures (align multi-layer mosaic option) available with TrakEM2 software (Cardona et al., 2012) (plug-in for NIH Image J, FIJI). In brief, for both within-layer and across-layer alignments, the expected transformation was set to Rigid to minimize scale change across layers while the desired transformation was set to Affine to minimize alignment error. Residual alignment jitter was further reduced by applying an Affine regularizer. Cell and circuit reconstructions were performed using TrakEM2 to first create skeletons of cones, bipolar cells, and ganglion cells. Terminal nodes within these skeletons were placed on synaptic ribbons so that the XYZ coordinates and the number of synapses could be determined for a sample of cells. Volume rendering of selected cell profiles and ribbon content (see Figure 1B) was performed manually by inspection of multiple annotators.

**Figure 1.**
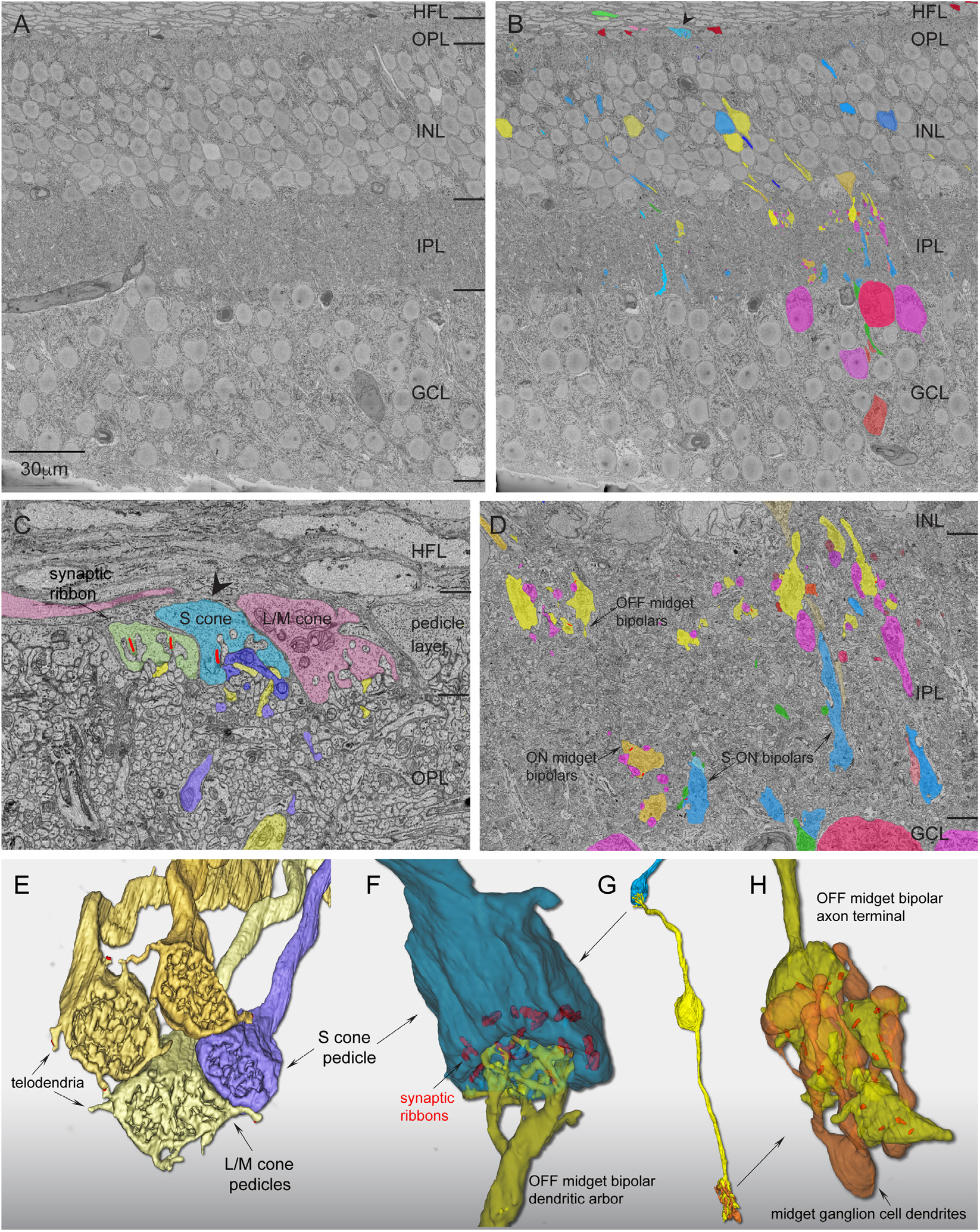
Scanning block-face electron microscopy demonstrates an S-OFF midget circuit in macaque retina. (A) Image of a single layer from the EM volume composed of a vertical section taken 400 μm from the foveal center along the foveal slope. At this central location, ganglion cell bodies stack up to 6 cell bodies thick. HFL, Henle fiber layer; OPL, outer plexiform layer; INL, inner nuclear layer; IPL, inner plexiform layer; GCL, ganglion cell layer. (B) Profiles of volume-rendered processes in another vertical layer. In the OPL, OFF midget bipolar profiles are in yellow and S-ON bipolar cells are in blue; at the top a pedicle identified as from an S cone is shown in light blue (arrowhead) alongside a few scattered rod pedicles (dark red). Below in the INL are OFF midget bipolar cells (yellow) and blue cone bipolar cells (blue). In the GCL, some midget ganglion cell bodies are shown in magenta and pink. (C) Zoomed view of three pedicles in the OPL: the S cone in panel B (blue), flanked by two L/M cones (green and red). Blue-cone bipolar dendritic arbors (dark blue profile) form the invaginating central element and are encircled by the flat dendritic arbors (yellow profiles) of an OFF midget bipolar cell. (D) Zoomed view of the IPL showing axons and axon terminals of OFF midget bipolar cells stratified near the outer border of the IPL (yellow profiles, arrow) and blue-cone bipolar cells stratified near the inner border of the IPL (blue profiles, arrows). A single ON midget bipolar terminal (orange profile; arrow) is also present. Midget ganglion cell dendritic profiles in the GCL are shown in magenta. (E) Complete reconstruction of four neighboring cone pedicles from one S cone and three L/M cones. The S cone pedicle is smaller than the L/M cones and lacks the distinctive telodendria that connect adjacent L/M cones. (F) Zoomed view of reconstructed S-cone pedicle in partial transparency, showing ribbon synapses (red) and the single dendritic arbor of an OFF midget bipolar in dense contact with the pedicle synaptic face. (G) Zoomed out view of the entire reconstructed OFF midget bipolar circuit (yellow) linked to the S-cone pedicle (blue) shown in F. The axon terminal exclusively contacts the compact dendritic arbor of an OFF midget ganglion cell (orange profile). (H) Zoomed view of the OFF midget bipolar axon terminal synaptic relation (yellow) with its associated OFF midget ganglion cell (orange) shown in partial transparency with synaptic ribbons (red). The ganglion cell for this particular private-line circuit was at the edge of the volume and could not be completely reconstructed.

### In vitro *preparation*

After retinal isolation as described above, radial cuts were made in the isolated retina-choroid to create a flat mount that was adhered, ganglion cell layer up, to the glass bottom of a thermostatically maintained (~36°C, TC-344B, Warner Instruments) steel superfusion chamber coated with poly-L-lysine (P1399, Sigma; 10 mg in 10 mL H_2_O). The retina was continuously superfused with Ames’ medium (pH 7.37; constant oxygenation with 95% O_2_/5% CO_2_; 3–5 ml min^−1^). Visual stimuli were projected onto the vitreal (ganglion-cell) side of the retina as *in situ*, via the microscope objective lens as described further below.

### In vitro *electrophysiology*

Retinal ganglion cells were observed using a 60× water immersion long working distance objective (1.0 NA, Nikon) under infrared illumination. Using the ‘loose-patch’ method, extracellular recordings were made with glass micropipettes (5–8 MΩ) filled with Ames’ medium. When possible, the recorded cells’ retinal location and distance from the foveal center was determined and recorded as temporal equivalent eccentricity (Watanabe and Rodieck, 1989). To map the cell’s receptive field, the cell body was first placed in the middle of the stimulus field.

Flashing white squares (2 Hz temporal frequency, 10 or 25 μm wide) were systemically moved in the *x* and *y* planes to locate the most sensitive point of the receptive field and determine its approximate size. The location of the maximum spike response was defined as the receptive-field midpoint; and visual stimuli were positioned relative to this point. Midget ganglion cells were visually identified and differentiated from other cell types by their relatively high density and small soma size (Dacey and Lee, 1994), which was confirmed by receptive-field mapping: midget ganglion cells showed the smallest center diameters of any primate ganglion cell (30–150 μm in the near retinal periphery). The other relatively high-density ganglion cell types (parasol and small bistratified cells) have relatively large cell bodies and thus show receptive field center diameters ~2–3 times larger than midget cells. Data acquisition and the delivery of visual stimuli were coordinated by custom software running on an Apple Macintosh computer. Current and spike waveforms were Bessel-filtered at 2 or 5 kHz and sampled at 10 kHz.

### Stimulus generation

A digital light projector (Christie Digital Systems) was used to project the visual stimuli (VSG, Cambridge Research Systems) through an optical relay to the microscope camera port and focus the image onto the photoreceptor layer. The irradiance spectra for red, green, and blue primaries were measured with a spectroradiometer (PR705, Photo Research, Inc.); peak wavelengths and integrated photon fluxes were 636, 550, and 465 nm and 2.7×10^6^, 6.9×10^5^, and 1.8×10^5^ photons s^−1^ μm^−2^, respectively. To compute the effectiveness of the light delivered by each primary to the cone aperture, we calculated the products of each primary irradiance spectrum and each cone spectral sensitivity function (Baylor et al., 1987). We corrected for the spectrally broadening effects of self-screening by assuming a pigment density of 0.016 μm^−1^ (Baylor et al., 1987) and a cone outer segment length of 5 μm: while cones at ~8 mm have an outer segment length of 20 μm (Banks et al., 1991), we correct for the fact that peripheral cones in vitro lay obliquely to the optical axis of the objective, thus shortening the effective path length and reducing light capture. Each product was then summed across wavelength giving units of ‘effective’ photons s^−1^ μm^−2^ (irradiance corrected by cone spectral sensitivity). Effective photons s^−1^ μm^−2^ were then converted to photoisomerizations s^−1^ cone^−1^ by multiplying by the area of the cone aperture. In previous studies involving transverse illumination of the cone outer segment (Baylor et al., 1979), where funneling of the inner segments plays no role, the conversion factor commonly used is 0.37 μm^2^. The efficiency of photoisomerization (0.67) (Dartnall, 1972) is included in this value. In the *in vitro* macaque retina, as *in vivo*, light is incident upon the vitreal surface of the retina and funneling by the inner segment would tend to increase the effective area of the cone aperture. We therefore consider the use of 0.37 μm^2^ a very conservative estimate of cone aperture to make the conversion to photoisomerizations s^−1^ cone^−1^.

Often, the intensity of stimuli used in human visual psychophysics or in physiological experiments in the intact primate eye is expressed in units of retinal illuminance, or Trolands (Td). To aid comparison with our data, we previously calculated that for a peripheral cone with an inner segment aperture of 9 μm, 1 Td was equivalent to ~30 photoisomerizations s^−1^ cone^−1^ (Crook et al., 2009).

To generate stimuli that isolated responses from only L, M, or S cones, we computed the linear transformation between the measured primary irradiance spectra and the known spectral sensitivity functions for L, M, and S cones (Baylor et al., 1987). The solution to this transformation outputs the mixture of red, green, and blue primary intensities required to evoke isolated responses from L, M, or S cones (Brainard and Stockman, 2010). Our cone-isolating stimuli have been previously confirmed by direct recordings from macaque cones (Packer et al., 2010). Cone contrast was defined as the peak excursion from the background, divided by the mean light level, expressed as a percentage. Computed contrast for cone-isolating stimuli was 18% around a background comprising equal quantal catches for the three cone types. Additional high-contrast S and L+M stimuli were 64% around an equal background.

### Identifying putative S-cone input and characterizing cone-specific contributions to the receptive field

Retinal ganglion cells were initially surveyed for putative S-cone inputs by recording extracellular spike activity while presenting spots of high-contrast S-cone or (L+M)-cone isolating stimuli to the cell’s receptive-field center (150 or 400 μm diameter, 64% contrast), modulated as square waves at 2 Hz temporal frequency. At this frequency, the ON response phase is ~0° and OFF response phase ~180°. Stimulus presentations were repeated 12–20 times and peristimulus time histograms (PSTHs) were constructed from the spikes. PSTHs were then fit with a Fourier series and spiking activity was reported as the amplitude, *A*, and phase, θ, of the first harmonic (F1). From this spiking activity, S or L+M mechanism-specific responses were classified as either ON or OFF with strength *R* = *A* cos(*θ*), where −*R* denotes an OFF response and +*R* denotes an ON response. While we always stimulated ganglion cells at photopic light levels to prevent against rod intrusion, this computation further penalizes any spurious rod response, which has a much longer latency to light increments (~270°; see Crook et al., 2009 Figure 5D). A minimum response threshold was determined by computing the response strength *R* for a subset of midget ganglion cells recorded during epochs of no stimulus presentation (1.00±0.62 spikes s^−1^, *n* = 56) and discarding any cell with response ≤4 SD from the mean (i.e., 3.48 spikes s^−1^). Cells with both |*R_L+M_*| ≤ 3.5 spikes s^−1^ and |*R_S_*| ≤ 3.5 spikes s^−1^ were considered unresponsive and were not included in further analyses. Midget cells with |*R_S_*| > 3.5 spikes s^−1^ were considered to have putative S-cone input.

To fully characterize the specific L-, M-, and Scone inputs to individual cells, extracellular spike activity was recorded while presenting full-field L-, M-, or S-cone isolating stimuli (800 or 1200 μm diameter, 18% contrast), modulated as square waves at a temporal frequency of 2 Hz. Stimulus presentations were repeated 12–20 times; PSTHs were constructed and fit with a Fourier series and spiking activity was reported as the amplitude, *A*, and phase, *θ*, of the first harmonic (F1). From this spiking activity, cone-specific responses were classified as either ON or OFF with strength *R* = *A* cos(*θ*), where −*R* denotes an OFF response and +*R* denotes an ON response. To compare the relative strength of L-, M-, and S-cone inputs to each cell, responses were normalized as 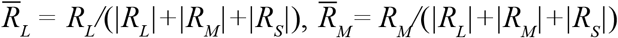, and 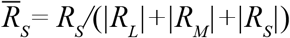. For midget cells when the initial high-contrast S or L+M stimulus was not run, we assessed responses to low-contrast stimuli with similar criteria as above to classify cells as unresponsive (|*R_L_*| ≤ 3.5 spikes s^−1^ and |*R_M_*| ≤ 3.5 spikes s^−1^).

### Computing spatial frequency tuning

The spatial tuning of midget-cell receptive fields was characterized using S cone–isolating or (L+M) cone–isolating drifting gratings (modulated at 2 Hz temporal frequency) of varying spatial frequency (spatial frequency 1/32–16 cpd, contrast 56% or 72%). For each cell, spike rate amplitude (*A*) and phase (*θ*) were computed from the first Fourier harmonic (F1) at each stimulus frequency, and tuning curves were constructed from pairs of amplitude/ phase measurements at each spatial frequency. Tuning curves were then fitted using a difference-of-Gaussians model (Enroth-Cugell et al., 1983) to determine the spatial arrangement of S and/or L+M inputs to the receptive field center and surround. Details and application of this model have been described previously (Dacey et al., 2000; McMahon et al., 2004; Wool et al., 2018). From the fitted tuning curves, the frequency of peak response, *f_peak_*, and the cutoff frequency, *f_cutoff_* (defined as the frequency at which response amplitude decreases to 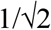 of the maximum) were computed for each cell for each stimulus. For a given cell, *f_cutoff_* > *f_peak_* indicates inputs are more localized to the smaller receptive-field center (i.e., tuned to higher spatial frequencies), while *f_cutoff_* < *f_peak_* indicates inputs are localized to the larger receptive-field surround (i.e., tuned to lower spatial frequencies).

### Computing three-dimensional color tuning of ganglion cells

To measure cells’ three-dimensional color tuning (i.e., beyond S, L−M, or L+M), we used a set of slow-modulating sinusoidal stimuli (Sun et al., 2006). These stimuli were constructed from a three-dimensional color space, *xyz*, defined by the three classical postreceptoral mechanisms of LGN neurons: a cone-subtractive (L−M) *x* axis, an S cone–isolating *y* axis, and a cone-additive (L+M) *z* axis (Derrington et al., 1984). When the chromaticity of a full-field stimulus is modulated around circles in the three planes formed by these axes, it can be decomposed into three phase-shifted cone-isolating sinusoids: in the isoluminant L−M versus S (*xy*) plane, L and M cones are in antiphase and are modulated 90° out of phase with S; in the L−M versus L+M (*xz*) plane, L and M are modulated 90° out of phase with each other; and in the S versus L+M (*yz*) plane, L and M are in phase but 90° out of phase with S. In any stimulus plane, vectors in the first and third quadrants represent additive input from the two mechanisms, whereas vectors in the second and fourth quadrants represent opponent inputs. Maximal cone contrast at the axes of the three stimulus planes was 8% L, 10% M, and 28% S (*xy*); 13% L, 14% M, and 0% S (*xz*); and 29% L, 29% M, 90% S (*yz*). The *x, y, z* notation used in this study denotes generic Cartesian coordinates in three-dimensional space, and should not be confused with the 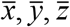 notation used in the CIE 1931 XYZ color space (Schanda, 2007)—another common color space but one we do not use in this study.

Each stimulus was presented as a uniform field (800–1200 μm) encompassing the cell’s receptive-field center and surround. Chromaticity was modulated at 1 Hz (360° s^−1^, or 6° per video frame) in a clockwise (CW) and a counterclockwise (CCW) direction, averaged to prevent response latency from biasing the time (and, thus, angle) of maximum response. Peristimulus time histograms (PSTHs, binwidth = 12°) for both directions were constructed from the average spike rate across 12–24 stimulus presentations. We performed Fourier analysis on each CW and CCW PSTH, and computed a response vector from the amplitude and phase of the first Fourier harmonic (F1). The cell’s preferred vector (*A, θ*) in a given plane was reported as the mean of the CW and CCW response vectors. To explore any clustering of preferred vectors in each stimulus plane, we used kernel density estimation to visualize the distributions of preferred vector angles, *θ* (preferred vector magnitude, *A*, was ignored) in response to each stimulus plane. For each cell type and stimulus plane, the distribution of preferred vector angles was binned (binwidth = 1°), convolved with a Gaussian filter (sigma = 10°), and then normalized.

The strength of each cell’s responsiveness to the L−M (*x*), S (*y*), and L+M (*z*) chromatic mechanisms was determined by first decomposing its preferred vector (*A, θ*) in each stimulus plane (*ISO: xy; LvM: xz; SvLM: yz*) into two axis components (*x* and *y, x* and *z*, or *y* and *z*) using Eqs. 1–6:

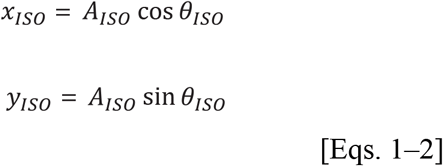

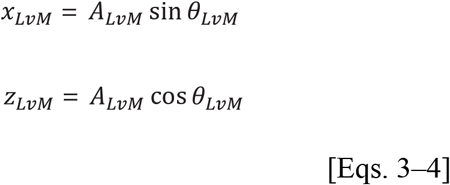

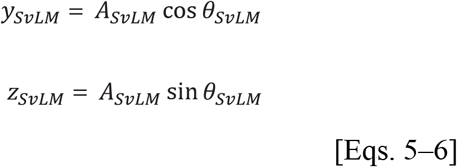

In these equations, *θ_ISO_, θ_LvM_* and *θ_SvLM_* are the preferred vector angles in the three planes, and *x, y*, and *z* values are the weights of each component in each two-dimensional plane. For each pair of components in Eqs. 1–6, effective contrast along stimulus axes was equalized by correcting one component by the contrast ratio between axes [e.g., *x_ISO_* × (*Cy_ISO_/Cx_ISO_*)]. Finally, to compute the [*X,Y,Z*] vector projection in three-dimensional color space (where *X* is the total contribution of the L−M mechanism, *Y* is the total contribution of the S mechanism, and *Z* is the total contribution of the L+M mechanism), each vector component was normalized by the total contrast of the stimulus plane, *C*, and preferred vector amplitude, *A*. Since each component is doubly represented across the three total stimuli, the weights of each component were averaged over the number of component measurements, *n*. For a cell where all three stimulus planes were presented (146 cells), *n* = 2 for each component, while for some cells where only two of the three total stimulus planes were presented (83 cells), *n* = 1 for two of the three components (Eqs. 7–9). We found computing a component’s weight to be reliable irrespective of one or two component measurements.

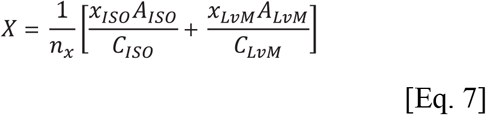

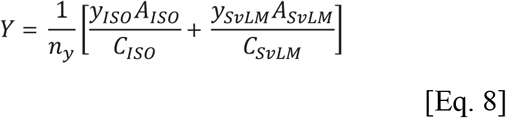

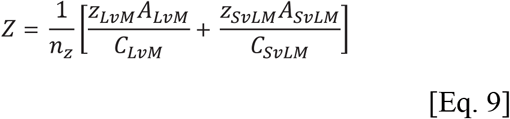

[*X,Y,Z*] coordinates were converted to spherical coordinates, and reported as the azimuth (*φ*, 0–360° on the *XY* isoluminant plane) and absolute value of the elevation (*θ*, 0–90°, where 0° falls on the *XY* plane and 90° falls orthogonal to it, along the *Z* axis).

### Statistics

All descriptive statistics denote mean±std unless otherwise noted, and directional statistics are employed to describe circular data (Berens, 2009). Two-sample Student’s *t* tests were used to compare L/M and S cone ribbon synapse numbers and to compare populations or distributions unless nonparametric data were assumed, in which case the two-sample Kolmogorov-Smirnov test was applied. All analyses were completed in MATLAB R2015b (The Mathworks).

## Results

### Connectomic reconstruction of parafoveal S-cone OFF midget pathway

We first identified putative S cones by known differences in pedicle morphology. The pedicles of S cones differ qualitatively from neighboring L and M cones being smaller in size and lacking the short telodendritic processes present in L and M cones that create a gap junction-coupled network across L and M cones (O’Brien et al., 2012). Figure 1 illustrates a single layer in our foveal volume and the location of a single pedicle identified as an S cone by these criteria (Fig. 1B–C, arrowhead). Figure 1E shows a reconstruction of this pedicle (blue) in relation to three neighboring, larger pedicles (yellow shades); each of the larger pedicles gave rise to distinct telodendria that made tip-to-tip contacts with each other. By contrast, the smaller putative S cone pedicle lacked telodendria and made no contacts with neighboring cone pedicles. This pedicle deployed 16 synaptic ribbons and was contacted by a single flat midget bipolar cell (Fig. 1D–G); this was reconstructed and all synaptic ribbons were localized and volume rendered within its axon terminal (Fig. 1G). This midget bipolar terminal contained 24 synaptic ribbons, all of which were in exclusive contact with a single midget ganglion cell dendritic tree (Fig. 1H). If this cone and others showing the same unique morphology are indeed S cones than they should form a selective ON pathway synaptic connection with the morphologically unique “blue-cone” bipolar cell type (Kouyama and Marshak, 1992). Moreover, these S cones should form a regularly spaced array (Shapiro et al., 1985; Curcio et al., 1991; Bumsted and Hendrickson, 1999) and comprise about 10% of the total number of cones in our volume.

There were 185 cone pedicles in the volume and we classified 17 of these cones (9%) as S cones based on their physical isolation from all larger neighboring cones; indeed, only these 17 pedicles lacked telodendria that contacted neighboring cones (Fig. 1E). There were also a number of rod spherules in our volume (*n* = 40; Fig 1B, top, dark red); consistent with a previous report, S cones—as well as L and M cones—gave rise to occasional telodendria that contacted the more outwardly situated rod spherules (O’Brien, et al., 2012). These telodendria were situated at the outer edge of the pedicle layer in alignment with the rod spherule layer and above the synaptic face of the cone pedicles where cone-to-cone telodendria are located. These pedicles also appeared to be dispersed regularly though the overall cone mosaic as is expected for S cones (Fig. 2A).

**Figure 2.**
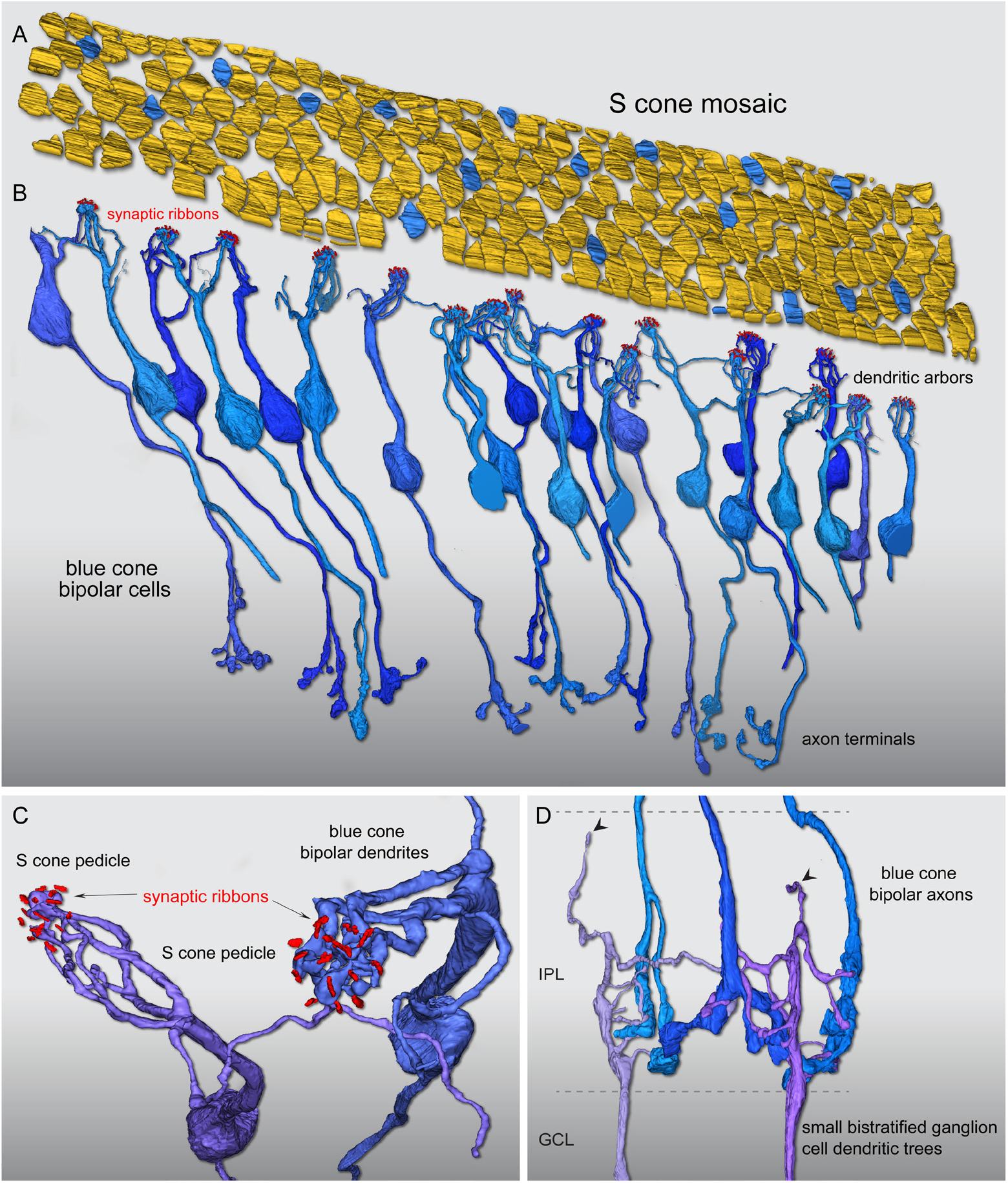
Reconstruction of foveal Scone mosaic and blue-cone bipolar cell synaptic pathway. (A) Outlines of the positions of 185 cone pedicle locations in the foveal volume (see Fig. 1). Seventeen regularly spaced cones (9%) were identified as S cones (blue) by their small size and lack of telodendritic contacts with neighboring larger pedicles (see Fig. 1). (B) Confirming the S-cone identity of these 17 pedicles, each received all invaginating central element contacts from a single cone bipolar type with the morphology of the blue-cone bipolar as previously described, with branching dendrites that converge on and selectively contact only S cone pedicles, and may contact 1–3 neighboring S cones. In this instance, we found 26 blue-cone bipolar cells in synaptic contact with these 17 S cones, occupying all central elements in the pedicle. (C) Zoomed view looking vertically through the pedicle of two blue-cone bipolar cell dendritic processes forming invaginating terminals (see also Fig. 1C) at neighboring S-cone pedicles. For visibility, the pedicle volume was made transparent; synaptic ribbon volumes (red) are shown in relation to the bipolar dendritic arbors. (D) The axons of the blue-on bipolar cells branch and terminate at the inner border of the IPL (see also Fig. 1D), where they make synaptic contact with presumed blue ON small bistratified ganglion cells. Two ganglion cell dendritic trees receive synaptic input from three blue-on bipolar cell axon terminals at the inner border of the IPL. These two ganglion cells also show very sparse dendrites that extend to the outer half of the IPL (arrowheads), consistent with their identification as the blue ON small bistratfied ganglion cell type.

We then reconstructed the bipolar cells linked to all invaginating central elements in each of these 17 cones. The result of this reconstruction (Fig. 2B) revealed that all central elements arose from a single cone bipolar type with the well-established morphology of the primate blue-cone bipolar cell (Mariani, 1984; Kouyama and Marshak, 1992). We found 26 blue-cone bipolar cells that formed divergent contacts exclusively to these 17 pedicles (1.5 bipolar cells per cone; Fig. 2B). A key identifying feature of this bipolar type is that it gives rise to several branching and occasionally long dendrites that bypass multiple cone pedicles to converge on one or two neighboring S cones (Fig. 2C) (Kouyama and Marshak, 1992).

Of the 26 bipolar cells whose dendritic arbors were reconstructed, 12 cells were completely contained in our volume and could be reconstructed to their axon terminals and synaptic contacts with ganglion cell dendrites in the inner plexiform layer (IPL) (Fig. 2B, D). Again, as expected for the blue cone bipolar cell type, the axonal arbor was branched and terminated at or near the inner border of the IPL. In addition, these axon terminals did not contact midget ganglion cells but instead made converging synaptic connections with a larger multibranched dendritic tree that gave rise to occasional branches that extended to the outer, OFF subdivision of the IPL. This is consistent with previous findings that the blue cone bipolar delivers ON-type S cone input to the inner dendrites of the small bistratified blue-ON ganglion cell type (Dacey and Lee, 1994; Calkins et al., 1998; Crook et al., 2009), which displays a bistratified dendritic tree.

Each of these 17 S cones also received extensive non-invaginating or basal contacts (Fig. 1C,F; Fig. 3; Fig. 4A), many of which were large and encircled the invaginating central element in the triad-associated position (Calkins, 2001; Tsukamoto and Omi, 2015). These large contacts arose exclusively from a single OFF midget bipolar cell for each of the 17 S cones in the foveal volume (Fig. 3A–B) identical to the pattern we observed for the first putatively identified S cone shown in Figure 1. These bipolar cells showed the distinctive midget bipolar dendritic and axonal morphology (Calkins et al., 1994): the cell body gave rise to a single stout dendrite that extended straight to the pedicle base before it arborized into a spray of short terminal branchlets at the synaptic face of the pedicle. This midget bipolar morphology contrasts with the relatively branched morphology of the blue cone bipolar cells (Fig. 2, Fig. 4B). Nine of these OFF midget bipolar cells could be reconstructed completely to their axon terminals in the outer portion of the IPL (Fig. 3C; see also Fig. 1D, H). The axon terminals formed a compact glomerular structure and private-line connection to an OFF midget ganglion cell classically described for foveal midget circuitry (Fig. 3D) (Kolb and Dekorver, 1991; Calkins et al., 1994).

**Figure 3.**
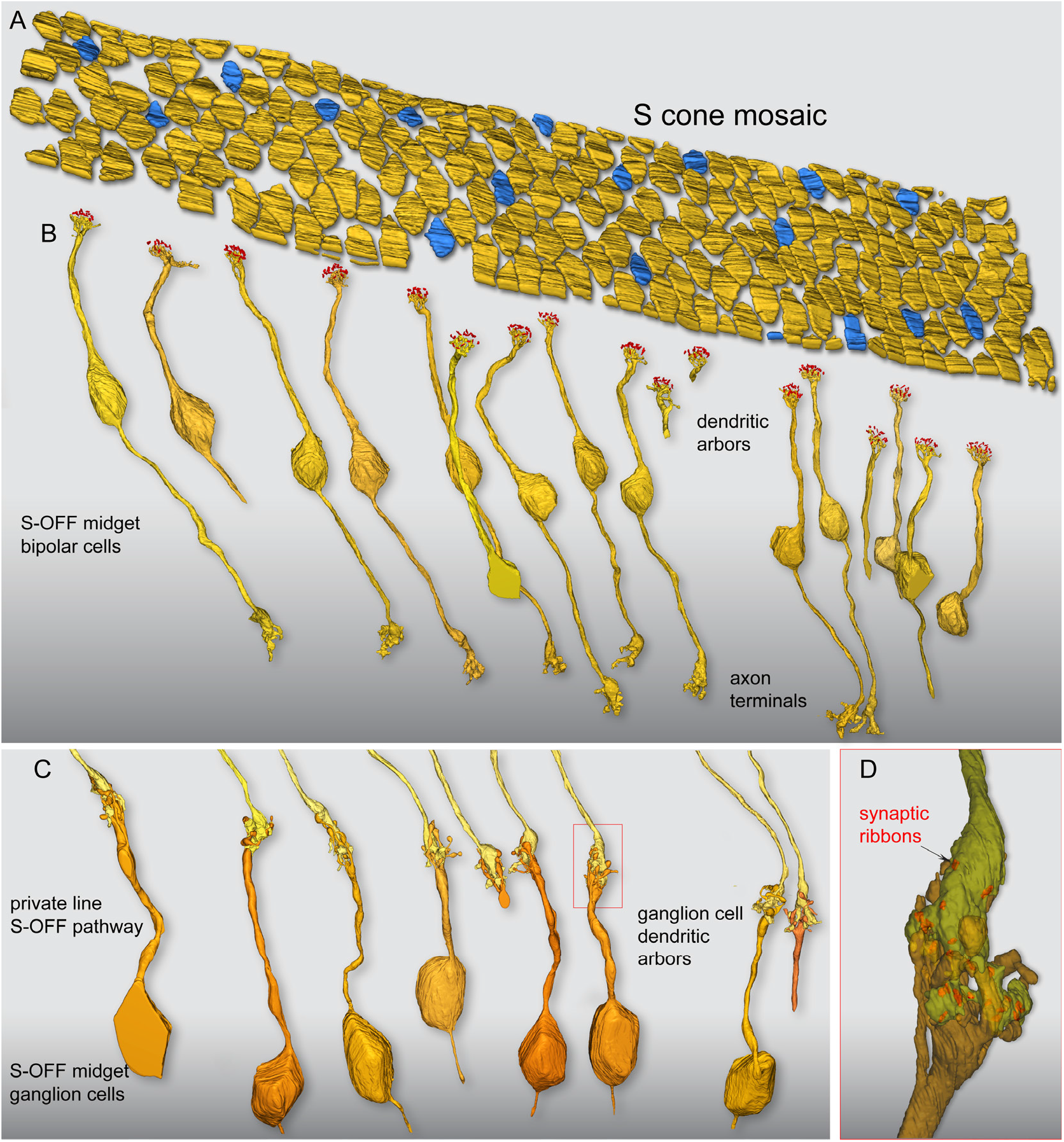
Flat contacts from midget bipolar cells form a private-line OFF pathway for S cones. (A) Outlines of pedicle positions with S-cone pedicles indicated in blue, as shown in Figure 2. (B) Each S-cone pedicle is densely contacted by the non-invaginating, ‘flat’ dendritic arbors of a single OFF midget bipolar cell. Of the 17 S-OFF midget bipolar dendritic arbors reconstructed, nine complete cells were encompassed within the volume and could be reconstructed completely to their axon terminals in the outer half of the IPL. (C) Axon terminals of the nine complete OFF midget bipolar cells shown in panel B; each forms a private-line synaptic connection with the dendritic arbor of an OFF midget ganglion cell. Seven of the nine midget ganglion cells could be reconstructed to the cell body and axon, which extends into the optic fiber layer. (D) One of the OFF midget bipolar–ganglion cell private-line synapses in panel C (red box), shown in greater detail. The S-OFF midget bipolar–ganglion cell private-line synaptic complex is shown in partial transparency; the bipolar terminal enters from above (yellow) and the ganglion cell dendritic arbor enters from below (orange). Twenty-five ribbon synapses in this bipolar cell were tagged and volume rendered (red), and established synaptic contact with this ganglion cell.

**Figure 4.**
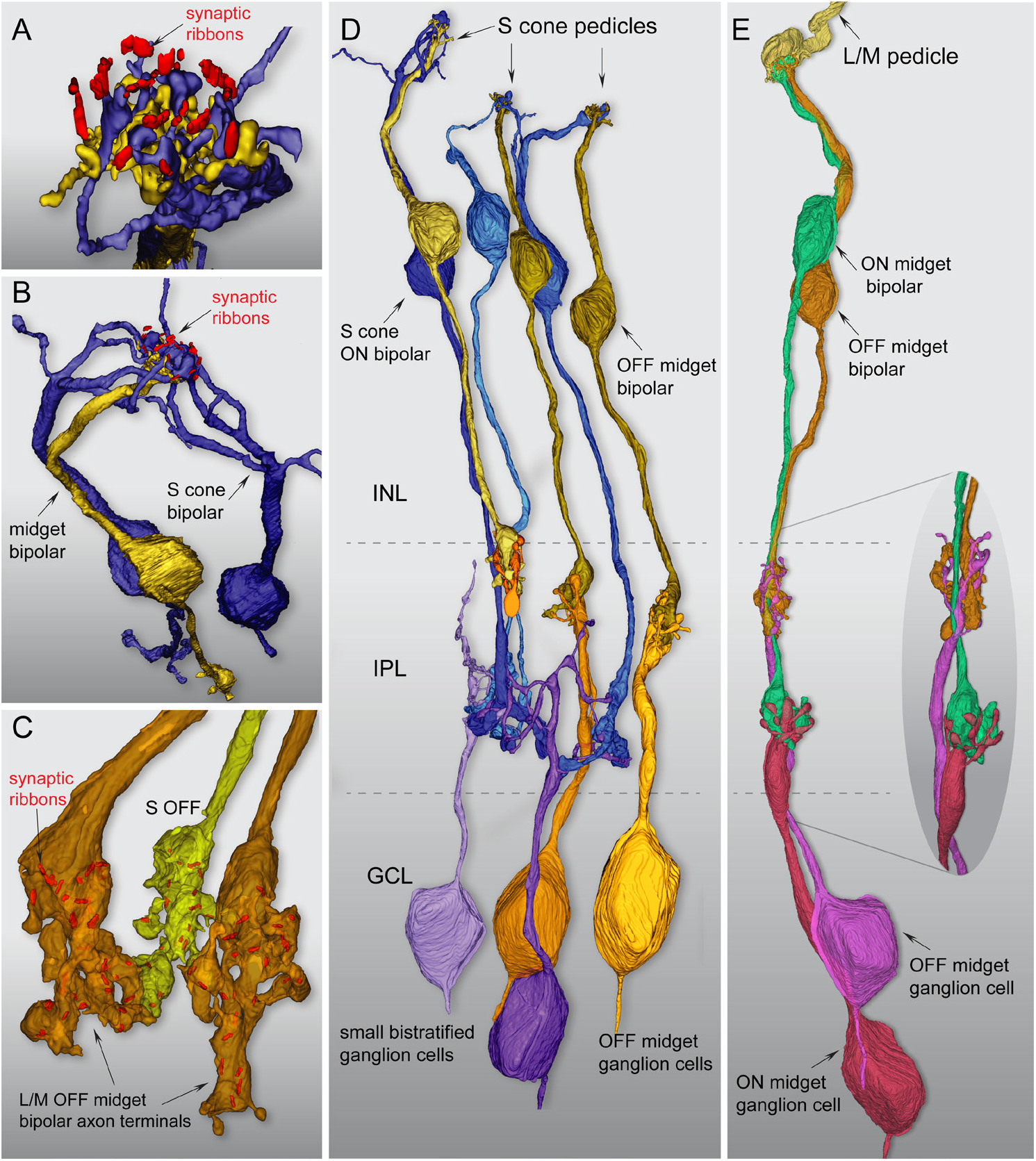
Summary of the morphological and synaptic asymmetry for ON and OFF S-cone pathways in macaque fovea. (A) Zoomed view looking down through an S-cone pedicle (transparent) at invaginating blue-cone bipolar (blue) and flat S-OFF midget bipolar (yellow) contacts, with the synaptic ribbon array (red). The blue-on bipolar cells give rise to very large terminations that occupy all central element positions in the pedicle, while the remaining fraction of the pedicle face is occupied by the flat contacts of a single OFF midget bipolar cell. (B) Another view looking down at a single S-cone ribbon array innervated by two branching and converging blue-cone bipolar dendritic trees (blue). In contrast, a single private-line connection of an OFF midget bipolar cell is present, in which a single stout dendrite forms a spray of terminal branches at the base of the pedicle (see also Fig. 1F). (C) Two L/M OFF midget bipolar axon terminals (orange) and a single neighboring S-OFF midget bipolar terminal (yellow), with synaptic ribbons shown in red. S-OFF midget bipolar cell axon terminals show the typical midget morphology but exhibit fewer synaptic ribbons relative to their L/M cone counterparts (see Results). (D,E) Fundamental difference in the ON and OFF circuitry originating from S cones and L/M cones in foveal retina. (D) On average, each S cone pedicle synapses with 1.5 blue-cone bipolar cells (blue) and 1 OFF midget bipolar cell (yellow). The blue-cone bipolar projects an axon terminal to the inner border of the IPL where it synapses with the blue ON small bistratified ganglion cell type (purple shades), while the OFF midget bipolar forms a private-line synapse with an OFF midget ganglion cell (yellow shades). (D) Reconstruction of a complete L/M cone midget circuit. The L/M cone pedicle (yellow) synapses with a single ON (green) and OFF (orange) midget bipolar cell; these in turn form private-line synaptic complexes with an inner-stratifying ON (red) and outer-stratifying OFF (magenta) midget ganglion cell, respectively. The entire circuit is entwined in close physical proximity throughout its length (inset).

The OFF midget bipolar cells that were associated with S cones differed quantitatively from their L/M cone counterparts in the number of synaptic ribbons present in the axon terminal (Fig. 3D, Fig. 4C). For the OFF midget bipolar cells connected to S cones the number of ribbon synapses (23.9±3.2, *n* = 9) was significantly lower than that found for a similar sample of neighboring OFF midget bipolar cells connected to L or M cones in the foveal volume (28.7±3.9, *n* = 9; *p* = 0.012, Student’s *t* test). Ribbon synapses were similarly reduced at the S cone pedicle relative to the L/M cone pedicles for these same cones with S-cone pedicles showing ~20% fewer synapse than their L/M cone counterparts; this difference was also significant (L/M cones = 21.67±1.00, *n* = 9; S cones = 16.89±1.7, *n* = 9; *p* = 0.0000018, Student’s *t* test). The parallel reduction in ribbon synapse numbers in both the S cone pedicle and the S-OFF midget bipolar terminal was not observed previously (Klug et al., 2003) and it was found that two S cone pedicles had a higher ribbon synapse number than the dominant L and M cone pedicles. Regardless, it does appear that the S-cone circuit can be identified by quantitative differences in synaptic architecture and that synaptic numbers in one part of the circuit can be mirrored at another point in the pathway.

In sum, we identified 17 S cones in our volume for the first time based on their unique pedicle morphology; these 17 cones also showed the expected density and spatial distribution expected for S cones. In addition, the central elements for these cones were all occupied by an ON bipolar type with the distinctive dendritic and axon morphology of the previously described blue cone bipolar cell (Mariani, 1984; Boycott and Wässle, 1991; Kouyama and Marshak, 1992; Herr et al., 2003). This further confirms their identity as S cones and establishes the unique convergence of morphological features that identifies the S cone circuitry in our foveal volume. We compared the connectivity observed at S cones to a sample of L/M cones in our volume (*n* = 17) and observed that ON midget bipolar cells occupied most of the central element positions at the pedicle. Each ON midget bipolar cell formed a private-line circuit with its ON midget ganglion cell partner and was tightly entwined with a private-line OFF midget bipolar circuit that mostly made triad-associated flat contacts with the pedicle (Fig. 4E), again as previously described (Kolb and Dekorver, 1991; Calkins et al., 1994). At the level single cell morphology, the L/M cone– and S cone–linked OFF midget circuits in this region appear identical. The presence of a private-line OFF midget bipolar–OFF midget ganglion cell connection associated with each of the 17 S cones clearly establishes this pathway as a major S-OFF pathway in the macaque monkey foveal retina. We thus sought to use physiological methods to functionally characterize this S-OFF midget signal, and determine its contribution to color tuning in midget ganglion cells outside of the foveal center where S, L and M cone inputs converge within the receptive field.

### Physiological targeting of midget ganglion cells in the near periphery

A main aim of this study was to amplify the impact of our structural characterization of an S-OFF midget circuit by also characterizing the functional properties of this circuit *in vitro*. As the midget pathway historically has been considered to accept inputs from only L and M cones, the S-cone connectivity that we observed in reconstructions of these cells should have a clear impact on how these cells respond *in situ* to complex visual stimuli—namely, color stimuli that combine signals from L, M, and S cones. To explore this, we recorded extracellular spiking activity from 404 retinal ganglion cells (28 blue ON, 281 OFF midget, 95 ON midget) in 37 retinae from male and female monkeys (*Macaca nemestrina, Macaca fascicularis*, or *Macaca mulatta*), targeting near-peripheral locations. As noted briefly in the Methods, midget ganglion cells in macaque retina could be unequivocally identified by their distinctive soma size, high maintained discharge rates, sustained light responses, and very small receptive-field center sizes (30-150 μm diameter) relative to all other ganglion cells; this is consistent with their very small dendritic tree diameters (Dacey and Petersen, 1992; Dacey, 1993; Diller et al., 2004; Crook et al., 2011; Wool et al., 2018). Parasol ganglion cells display large cell body diameters, transient light responses, and receptive fields ~3 times the diameter of midget cells. All other ganglion cell types exhibit receptive fields much larger than that of parasol cells (Crook et al., 2008). The distinction between midget ganglion cells and blue ON cells is also clear due to the blue ON cell’s distinctive response to S-cone stimuli and its large receptive field size (Crook et al., 2009), as considered further below. As inputs to midget ganglion cells increase as a function of eccentricity (Wool et al., 2018), cells in our recording locations sample anywhere from 3 to 12 cones, theoretically maximizing the likelihood of encountering an S cone in a cell’s receptive field. Receptive-field mapping (see Methods) was used to differentiate midget ganglion cells from their larger (2–3×) blue ON counterparts. We then used a range of stimuli to characterize each cell’s cone-specific inputs, spatial properties, and chromatic signature. While our aim was to record all cells’ responses to the full set of stimuli, this was not always possible; thus partial characterizations are also reported and the number of cells recorded during each stimulus is indicated. When possible, we recorded each cell’s retinal location in units of temporal equivalent eccentricity. Cells were located on the range of 9–48° (30±7°, *n* = 331).

### Some OFF midget ganglion cells respond to S-cone stimulation

We delivered high-contrast S-cone or (L+M)-cone isolating stimuli (spots of 150 or 400 μm diameter, 64% contrast) to each cell’s receptive-field center. While initial receptive-field mapping easily distinguished blue ON cells from midget ganglion cells due to their relatively larger receptive-field size, blue ON cells were moreover characterized by a vigorous ON response to S-cone stimulation. To these square-wave stimuli, responses to S-cone stimulation was entirely absent in most ON and OFF midget cells, and these cells were instead more reliably characterized by the phase of their response to (L+M) stimulation: ON cells responded in the ON phase (0–250 ms, or ~0°) of the stimulus (Fig. 5A), while OFF cells responded in the OFF phase (250–500 ms, or ~180°; Fig. 5B). In comparison, all blue ON cells responded to S-cone stimulation in the ON phase, and responded to (L+M)-cone stimulation in the OFF phase (Fig. 5C). Notably, in a small subset of OFF midget cells, the characteristic OFF response to (L+M)-cone stimulation was accompanied by a small response to S-cone stimulation in the same phase (Fig 5D).

**Figure 5.**
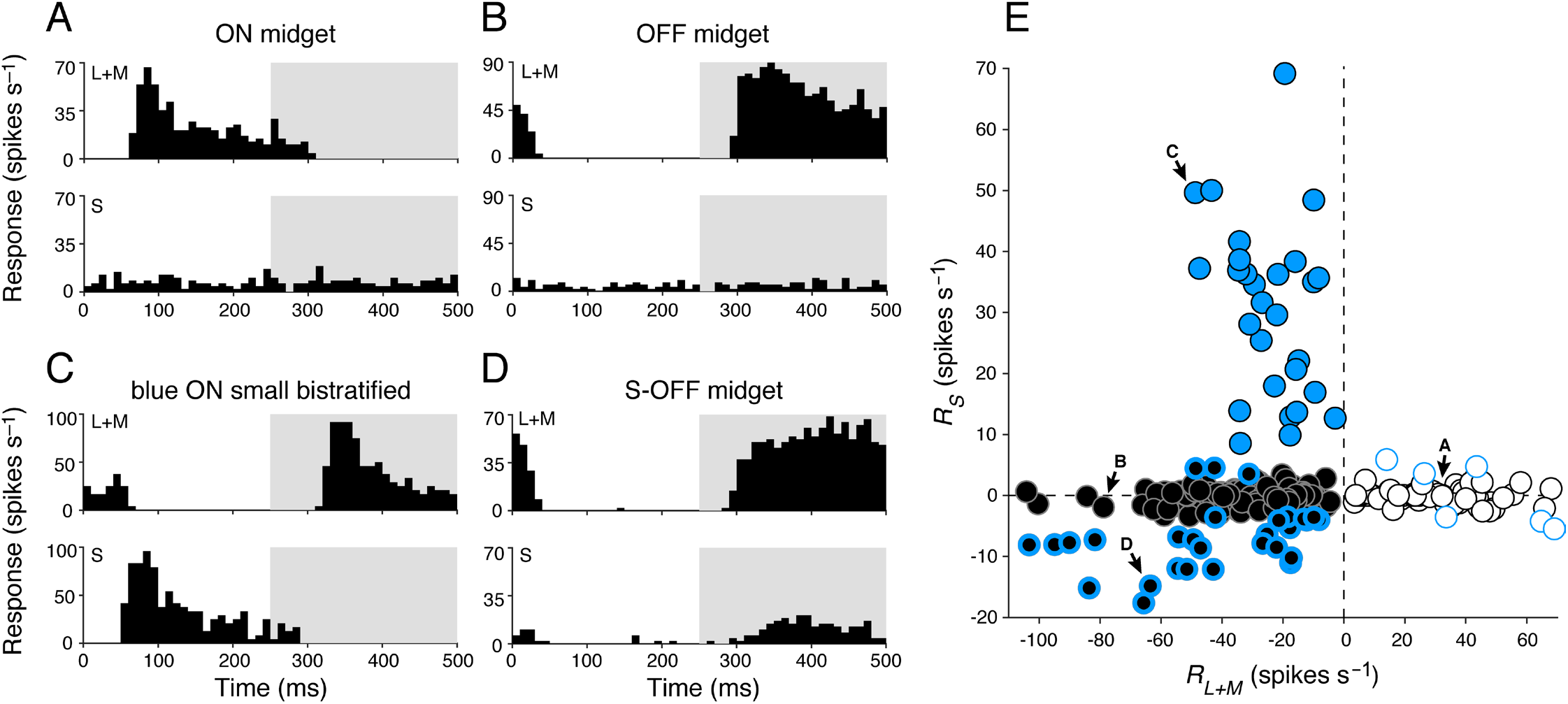
Cone-mechanism-specific responses in retinal ganglion cell subtypes. (A) Peristimulus time histogram of a typical ON midget ganglion cell shows spiking activity in the ON phase (0–250 ms) of a 2-Hz modulating (L+M)-cone-isolating stimulus (top) while showing no activity during a modulating S-cone-isolating stimulus (bottom). (B) A typical OFF midget cell shows the reverse behavior, with spiking activity in the OFF phase (250–500 ms) of a 2-Hz modulating (L+M)-cone-isolating stimulus (top) but also showing no activity during a modulating S-cone-isolating stimulus (bottom). (C) A blue ON ganglion cell responds to both L+M (top) and S (bottom) cone-isolating stimuli, with opposite phase. (D) An S-OFF midget ganglion cell demonstrates classical sensitivity to (L+M)-cone stimuli (top), as well as a small response to S-cone stimuli (bottom); both responses occur in the OFF phase. (E) Ganglion cell types cluster based on their responsiveness to S- (*R_S_*) and (L+M)- (*R_L+M_*) cone stimulation. Blue circles, blue ON cells; white circles, ON midget cells; black circles, OFF midget cells. Blue-bordered white and black circles denote midget ganglion cells with |*R_S_*| > 3.5 spikes s^−1^. Labeled cells are those shown in panels A–D, respectively.

To compare the relationship of S- and (L+M)-cone response properties of blue ON, ON midget, and OFF midget ganglion cells (including any midget ganglion cells with putative S-cone input), component mechanism–specific responses *R_S_* and *R_L+M_* were computed from the amplitude (*A*) and phase (*θ*) of spike activity during S and (L+M) stimulation (see Methods) to determine the relative strength of each mechanism. Blue ON, OFF midget, and ON midget cells form distinct clusters based on their relative responsiveness to S- or (L+M)-cone stimulation (Fig. 5E). ON and OFF midget cells are well defined by the sign of response to L+M stimulation (OFF: *R_L+M_* = −28.4±16.9 spikes s^−1^, ON: *R_L+M_* = 24.9±16.1 spikes s^−1^), and most cells’ lack of response to S-cone stimulation caused clustering at the *x* axis (OFF: *R_S_* = −1.1±0.90 spikes s^−1^, ON: *R_S_* = 0.51±0.83 spikes s^−1^). Blue ON cells showed balanced and opponent S-ON and (L+M)-OFF responses (*R_S_* = 30.5±14.5 spikes s^−1^, *R_L+M_* = −24.0±12.0 spikes s^−1^). While most ON and OFF midget cells showed no S-cone response, the small subpopulation that did (OFF cells, in particular) fall farther off the *x* axis. We classified as putative S-cone midgets those cells with |*R_S_*| > 3.5 spikes s^−1^, which was <4 SD above the mean *R* for a subset of cells where responses were recorded during epochs of no stimulus presentation (1.00±0.62, *n* = 56). This criterion identified a subset of 35 S-cone midget cells (29 OFF midgets, 6 ON midgets). For the majority of cells in this subset (26 OFF midgets, 3 ON midgets), responses to S-cone stimulation occurred in the same phase as responses to L+M stimulation (OFF: *R_S_* = −8.2±4.0 spikes s^−1^, *R_L+M_* = −43.2±29.1 spikes s^−1^; ON: *R_S_* = 4.7±1.1 spikes s^−1^, *R_L+M_* = 28.2±14.8 spikes s^−1^). The remaining cells (3 OFF midgets, 3 ON midgets) showed S-cone responses in the opposite phase.

### Spatial tuning of S-OFF midget cells identifies S-cone input to receptive-field centers

We next sought to identify the arrangement of S-cone input to the receptive fields of S-OFF midget cells. All ON and OFF midget ganglion cells exhibit classical center-surround receptive-field structure: a small central ON- or OFF-responsive region (formed by excitatory inputs from midget bipolar cells), surrounded by a broader region of opposite-phase sensitivity (formed by inhibitory inputs from horizontal cells) (Crook et al., 2011). Thus, the OFF-phase S-cone response we observed in OFF-center midget cells is consistent with an anatomical arrangement where S-cone input is propagated by an OFF midget bipolar cell that contributes to the receptive-field center. We further explored this hypothesis by characterizing cells’ spatial properties to understand whether S-cone input was localized to the smaller receptive-field center (i.e., sensitive to higher spatial frequencies) or to the larger receptive-field surround (i.e., sensitive to lower spatial frequencies). For 145 midget ganglion cells (114 OFF midgets, 12 ON midgets, 19 S-OFF midgets) we measured spike discharges to (L+M)-cone and/or S-cone drifting gratings of increasing spatial frequency. While S-OFF cells’ sensitivity to S-cone stimulation encourages an initial comparison to blue ON cells, the retina’s more well-known substrate for S-cone signals, we found the spatial tuning of S-OFF midgets to be qualitatively different from the spatial tuning of blue ON cells, suggesting a functional asymmetry in how S-cone signals are transmitted across the two ganglion-cell subclasses (Fig. 6). While blue ON cells show low-pass tuning to both (L+M)- and S-cone gratings with a clear antiphase relationship between mechanisms (this is due to spatially coextensive receptive fields; see Crook et al., 2009 Fig. 6A), the responses of S-OFF midget cells showed different spatial tuning and phase properties during (L+M)- and S-cone gratings (Fig. 6B), with a spatial profile much more similar to other OFF midget cells.

**Figure 6.**
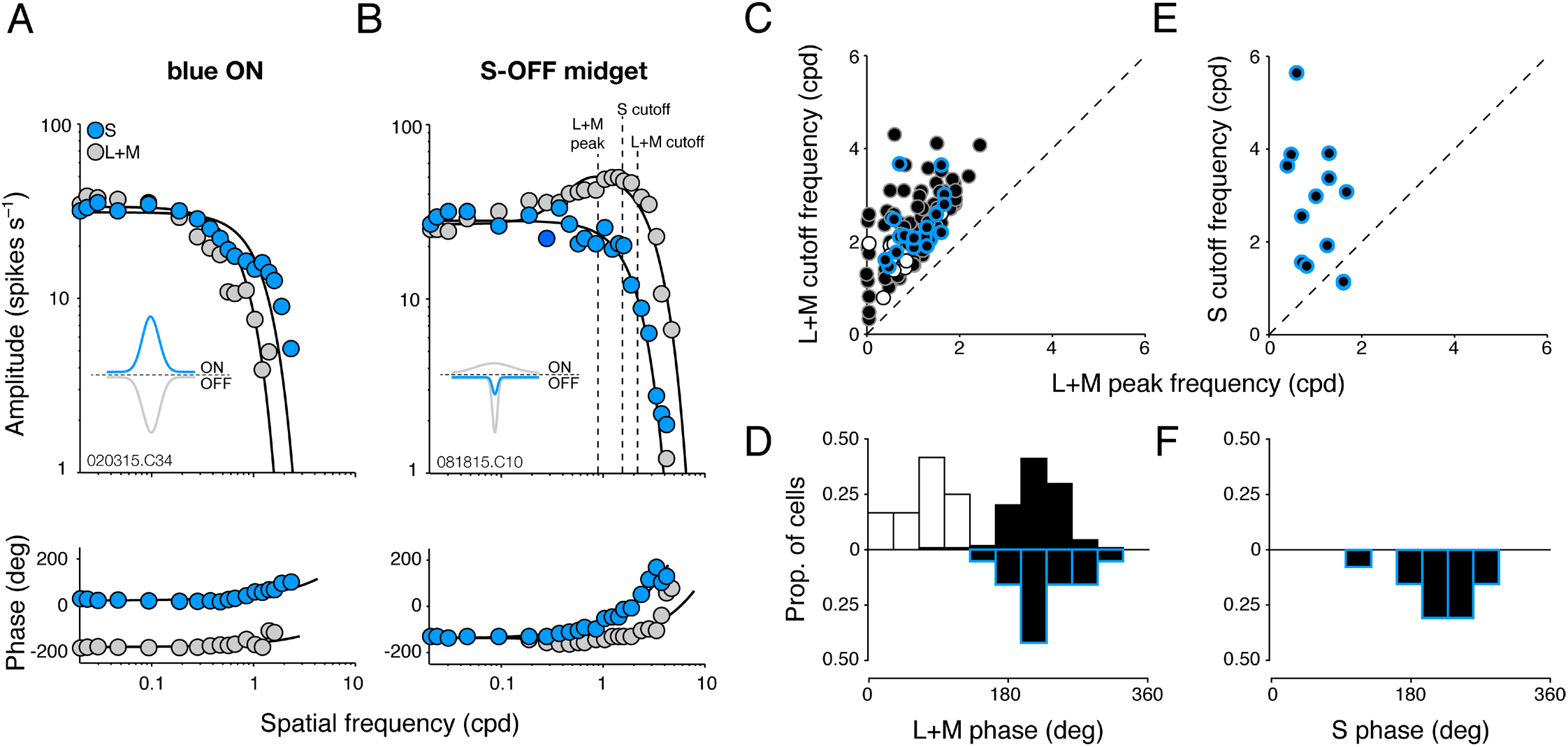
Spatial extent of S-cone input to OFF midget ganglion cells. (A,B) Compared to blue ON ganglion cells, which demonstrate spatially coextensive, low-pass, and antiphase spatial tuning to both S- and (L+M)-cone gratings (A), S-OFF midget ganglion cells demonstrate a low-pass spatial sensitivity to S-cone gratings atop the typical band-pass sensitivity to (L+M)-cone gratings, which occurs in the same phase (B). Inset Gaussian curves illustrate the type of receptive-field profile expected from each cell. (C) All midget ganglion cells demonstrated characteristic band-pass tuning to L+M gratings, in which the cutoff frequency for L+M gratings occurs at a higher spatial frequency than the peak frequency. The L+M spatial profiles of putative S-OFF midget ganglion cells (black/blue circles) are compared to typical ON (white circles) and OFF (black circles) midget cells, with cutoff frequency for L+M gratings (y axis) plotted as a function of peak frequency (x axis). (D) Distribution of response phase to L+M gratings for S-OFF (black/blue bars), ON (white bars), and OFF (black bars) midget cells. (E) For S-OFF midget cells, cutoff frequency for S gratings (y axis) is plotted as a function of peak frequency for L+M gratings (x axis). (F) Distribution of response phase to S gratings, for S-OFF midget cells.

In response to L+M drifting gratings, all OFF, ON, and S-OFF midget cells showed characteristic band-pass tuning (diminished and antagonistic center-surround responses at low spatial frequencies, and enhanced center-isolating responses at high spatial frequencies; Fig. 6B). As such, there was a clear *f_peak_* for each cell and *f_cutoff_* > *f_peak_* for all cells (Fig. 6C). For OFF midget cells without S-cone input, *f_peak_* = 1.02±0.56 cycles degree^−1^ (cpd) and *f_uutoff_* = 2.23±0.75 cpd. For ON midget cells, *f_peak_* = 0.65±0.37 cpd and *f_cutoff_* = 1.65±0.44 cpd. The slightly lower spatial tuning of ON midget cells has been previously described, and is attributed to slightly larger dendritic trees (Watanabe and Rodieck, 1989; Dacey and Petersen, 1992; Dacey, 1993). By comparison, S-OFF midget cells showed a similar spatial profile to other OFF midget cells in response to L+M drifting gratings, with *f_peak_* = 1.08±0.43 cpd and *f_cutoff_* = 2.30±0.61 cpd. Neither *f_peak_* nor *f_cutoff_* was statistically significantly different between OFF midget cells with or without S-cone input (*p* = 0.65 and *p* = 0.73, respectively; Student’s *t* test). As expected, the phase of response to L+M drifting gratings between ON and OFF midget cells differed by ~180° (ON = 75.5±34.8°, OFF = 216.2±33.2°). In S-OFF midget cells, we found that the phase of response to L+M drifting gratings (224.8±40.9°) closely matched that observed in the other OFF midget cells; no statistically significant difference of response phase was found between the two populations (*p* = 0.31; Student’s *t* test) (Fig. 6D).

While the spatial tuning and phase of L+M inputs to S-OFF midget cells closely resemble those of typical OFF midget cells, we also characterized the spatial tuning and phase properties of their S-cone sensitivity using S-cone drifting gratings (14 S-OFF cells). In contrast to the band-pass tuning curves observed for L+M drifting gratings, responses to S-cone drifting gratings showed low-pass spatial tuning and were thus best fit with a single Gaussian function. The cutoff frequency of most S-OFF cells during S-cone stimulation (*f_cutoff_* = 2.79±1.33 cpd) was greater than the peak frequency as identified during L+M stimulation (Fig. 6E); this high spatial frequency tuning suggests an arrangement of S-cone input sequestered to the receptive-field center. This is further supported by the phase of response to S-cone drifting gratings we observed in these cells (218.5±43.9°), which closely matches the response phase of both S-OFF and OFF midget cells during presentation of L+M drifting gratings. This S-cone spatial profile of S-OFF cells that we describe—low-pass tuning, high *f_cutoff_*, and a response phase similar to other OFF cells—suggests that the S-cone contribution we observed in these cells is localized to the receptive field center, with a very weak and/or infrequent contribution from the receptive-field surround. This is consistent with the bipolar cell-mediated S-cone connectivity we observed anatomically, which predicts functional inputs localized in this way.

**Figure 7.**
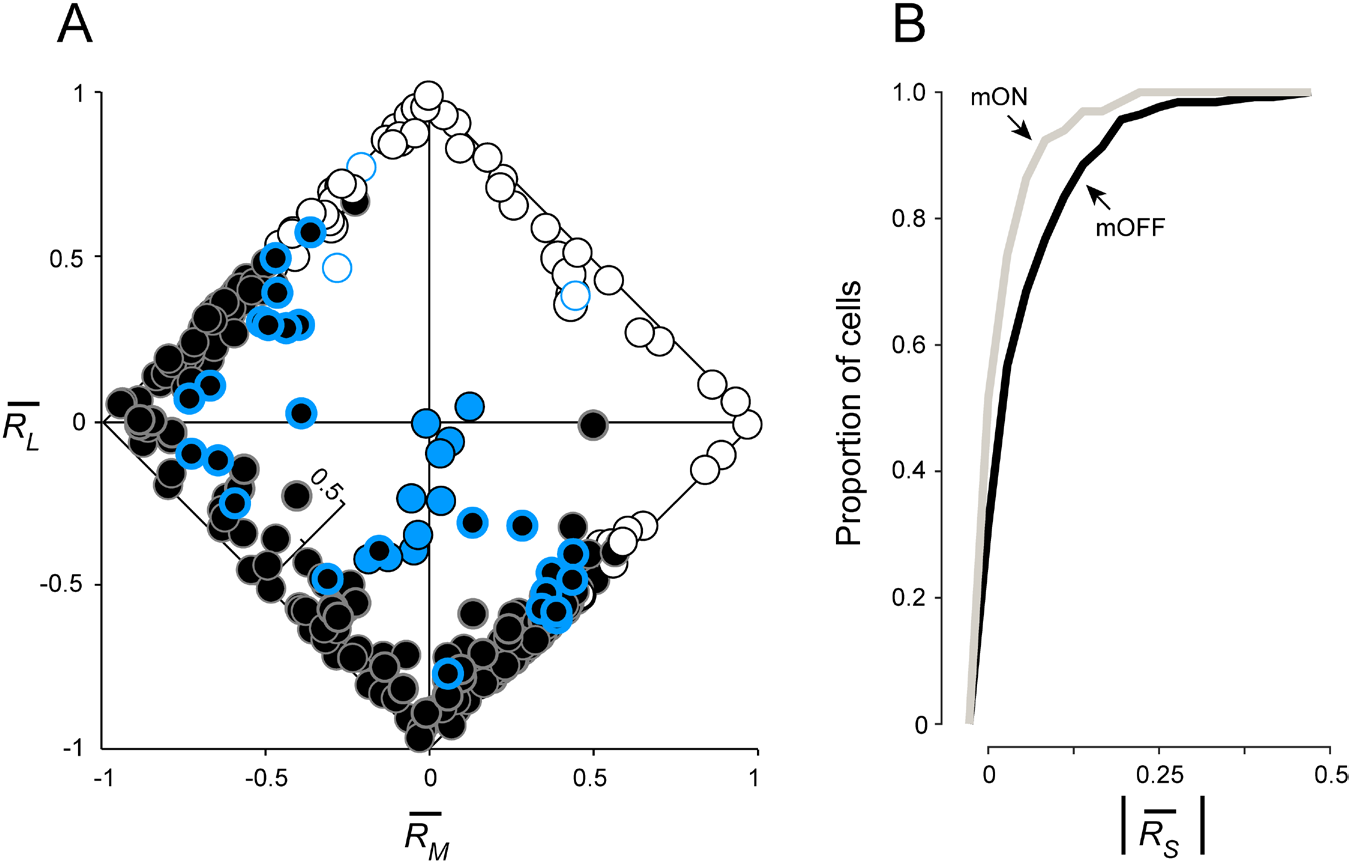
L-, M-, and S-cone-specific inputs to ganglion-cell subtypes. (A) Cone-specific responses *R_L_, R_M_*, and *R_S_* are shown for each ganglion cell by plotting relative L-versus M-cone input on the *x* and *y* axes, respectively. S-cone input is denoted by a third internal *z* axis. Classical L−M opponency is assessed by comparing the sign of *R_L_* and *R_M_* opponent cells fall along the diagonals where the signs of *R_L_* and *R_M_* are opposed, while nonopponent cells fall a long the diagonals where the signs are matched. Blue ON cells (blue circles) demonstrate sign-matched L- and M-cone inputs, along with strong S-cone input. ON (white circles) and OFF (black circles) midget cells demonstrate both opponent and nonopponent L- and M-cone inputs, with OFF midget cells demonstrating stronger S-cone input overall. Blue-bordered white and black circles denote ON and OFF midget cells initially identified as S-cone-sensitive in Figure 1. (B) Cumulative distribution of proportional S-cone input to OFF midget ganglion cells (black) compared to ON midget ganglion cells (gray).

### S cones contribute to both color-opponent and nonopponent OFF midget cells

After identifying a subgroup of S-OFF midget cells and characterizing the spatial properties of their S-cone input, we then asked how this input was related to the classical L−M color opponency observed in the midget pathway. We thus sought to determine whether the Scone response had any systematic relation to the sign of either the L or M cone input to the receptive field. To answer this, we recorded extracellular spikes from 332 ganglion cells (10 blue ON cells, 256 OFF midget cells, 66 ON midget cells) during full-field L-, M-, and S-cone-isolating stimulation with square-wave stimuli (18% contrast, 800 or 1200 μm diameter), then computed the component mechanism-specific responses *R_L_, R_M_* and *R_S_* from each cell’s F1 amplitude (*A*) and phase (*θ*). To compare the proportional strength of L-, M-, and S-cone inputs to each cell, responses were normalized as 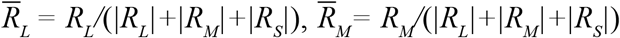, and 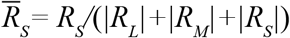.

Midget cells were characterized as cone-opponent or nonopponent by comparing the sign of 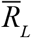 and 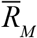. ON and OFF midget cells were considered opponent when L- and M-cone responses differed in sign (e.g., 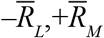 or 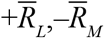), and nonopponent when responses had the same sign (e.g., 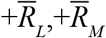 or 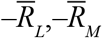). Consistent with our previous recordings in near-peripheral retina (Wool, 2018), our midget cell population was heterogeneously composed of both L−M opponent (195 OFF, 44 ON) and nonopponent (61 OFF, 22 ON) cells (Fig. 7A). Comparing the relative unsigned proportion of S-, L-, and M-cone inputs to each cell, we observed a distinct cluster of blue ON cells in which S-cone input comprised the majority of cone input (0.70±0.19), against nonopponent L- and M-cone inputs. By comparison, we found that midget cells formed a broad continuum of L−M opponency, but also of S-cone sensitivity, particularly in OFF midget cells. The mean proportional S-cone input across ON midget cells was 0.04±0.05, while across OFF midget cells the proportion was twice as high, 0.08±0.09. The stronger overall S-cone contribution to OFF midget receptive fields is visualized in Figure 7A as a greater scatter of OFF midget cells into the orthogonal axis; ON midget cells, by contrast, remain more closely clustered along the diagonals between the dominant L and M axes. To further quantify this tendency of OFF midget cells to exhibit stronger S-cone input than ON midget cells, we compared the cumulative distributions of ON and OFF midget cells as a function of unsigned proportional S-cone input, (Fig. 7B). ON midget cells show a narrower distribution around weak or absent S-cone input, while S-cone input in OFF midget cells is more broadly distributed toward higher values of. This distribution of S-cone input between ON and OFF midget cells was statistically significantly different (*p* = 0.0013, Kolmogorov-Smirnov test). Indeed, a much larger percentage of OFF midget cells showed proportional S-cone input of >0.10: 27.3% (70/256), compared to 10.6% (7/66) of ON midget cells.

Nonopponent S-OFF midget cells were much more prevalent in our sample compared to opponent S-OFF cells. OFF midget cells with proportional S-cone input of >0.10 comprised 45.9% (28/61) of all nonopponent OFF midget cells, but only 21.5% (42/195) of opponent OFF cells. Since the likelihood of encountering opponent or nonopponent midget cells is itself a function of eccentricity (Wool et al., 2018), we assessed whether this difference in detectable S-cone input between opponent and nonopponent OFF midget cells could also be attributed to a difference in retinal location. We found the mean eccentricity between these groups to be statistically indistinguishable (nonopponent cells: 33.3±6.6°, *n* = 24; opponent cells: 30.7±6.8°, *n* = 34; *p* = 0.15, Student’s *t* test). While we did not further test whether nonopponent OFF midget cells have a greater likelihood of exhibiting S-cone input, the effect at least appears to be independent of eccentricity-dependent receptive-field sampling of the cone mosaic. In any case, our analysis of cone-specific inputs to midget cells demonstrated that a substantial number of both opponent and nonopponent OFF midget cells exhibit not only classical L- and M-cone inputs, but in fact input from all three cone types. As the degree of L, M, and S sensitivity manifests heterogeneously across this class of S cone–sensitive midget ganglion cell, we sought to further probe these interactions by assigning a precise chromatic signature to each cell, identifying where these cells would fall in a complex, three-dimensional color space.

### S-OFF midget cells demonstrate complex, three-dimensional color tuning

A major goal of this study was to determine how the three classical postreceptoral color mechanisms (as identified in LGN neurons) (Derrington et al., 1984) combine at the level of the midget ganglion-cell receptive field to encode color sensitivity beyond L−M (‘red-green’) opponency. In a three-dimensional color space, *xyz*, defined by a cone-subtractive (L−M, *x*) axis, an Scone isolating (*y*) axis, and a cone-additive (L+M, *z*) axis, classical ‘red-green’ opponent midget cells would simply project along the single L−M dimension. While we observed many cells with such classical opponency, we also observed many midget ganglion cells with unbalanced L and M inputs, nonopponent L and M inputs, or even S-cone inputs (Fig. 7A). We thus expected the color preferences of these cells to be heterogeneous and to project away from the single L−M opponent axis.

To assess the strength of the three postreceptoral mechanisms in individual ganglion cells, and visualize their color preferences in this three-dimensional color space, we used a set of slow-modulating sinusoidal stimuli (Sun et al., 2006) in which chromaticity was modulated around circles in three intersecting stimulus planes: an (L−M) versus S isoluminant plane (*xy*, Fig. 8A), an (L−M) versus (L+M) plane (*yz*, Fig. 8B), and an (L+M) versus S plane (*xz*, Fig. 8C). Each stimulus was presented as a uniform field (800–1200 μm) encompassing the cell’s receptive-field center and surround. For an example classical OFF midget cell with balanced and opponent L and M inputs and no S input, PSTHs show the cell’s preference for −L+M stimuli in each of the three planes: near 180° (−L+M) in the *xy* plane (Fig. 8D), near 135° (+M–L) in the *yz* plane (Figure 8E), and near 270° (−L−M) in the *xz* plane (Fig. 8F). By contrast, an example opponent OFF midget cell with S-cone input shows significantly shifted preferred vectors that reflect not only the cell’s preference for opponent −M+L stimuli, but also additional S-cone sensitivity to its receptive field: near 315° (−S, +L, −M) in the *xy* plane (Fig. 8G), near 0° (+L, −M) in the *yz* plane (Fig. 8H), and near 180° (−S) in the *xz* plane (Fig. 8I).

**Figure 8.**
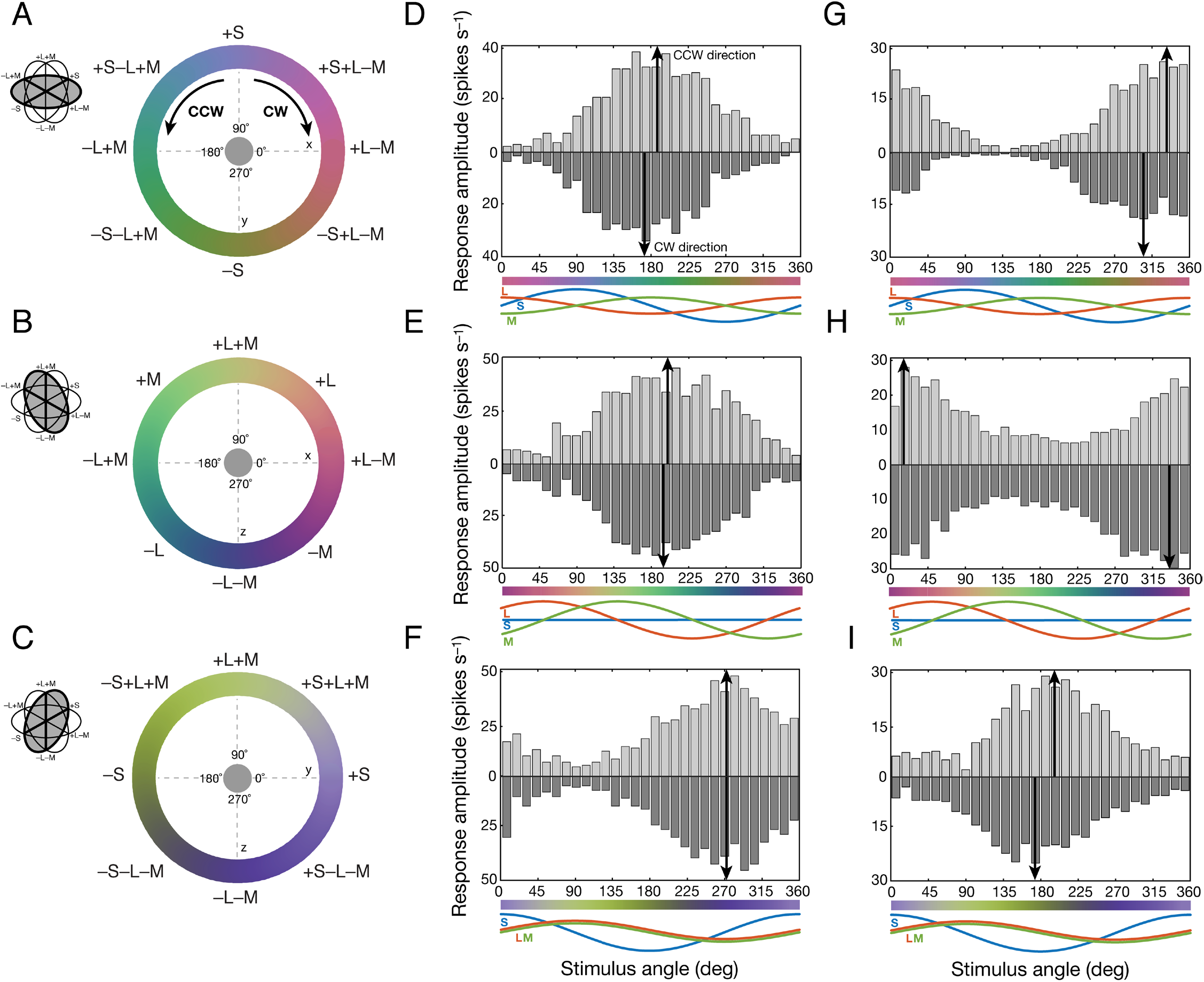
Measuring color preferences of individual ganglion cells. (A–C) The chromaticity of a uniform, full-field stimulus was modulated in clockwise (CW) or counterclockwise (CCW) directions along around circles in three intersecting planes of three-dimensional color space, xyz: an (L−M) versus S isoluminant plane (*xy*, A), an (L−M) versus (L+M) plane (*yz*, B), and an (L+M) versus S plane (xz, C). (D–F) Peristimulus time histograms reveal the color preference in each stimulus plane for an L cone–dominated, opponent OFF midget ganglion cell with absent S-cone input (61% L, 39% M, 0% S). The cell peaks near 180° (−L+M) in the *xy* plane (D), near 135° (+M−L) in the *yz* plane (E), and near 270° (−L−M) in the *xz* plane (F). (G–I) By contrast, peristimulus time histograms reveal the color preference in each stimulus plane for an M cone-dominated, opponent S–OFF midget ganglion cell with inputs from all three cones types (23% L, 44% M, 33% S). This cell peaks near 315° (−S, +L, −M) in the *xy* plane (G), near 0° (+L, −M) in the *yz* plane (H), and near 180° (−S) in the *xz* plane (I).

We recorded spike activity from 223 retinal ganglion cells (10 blue ON cells, 191 OFF midget cells, 22 ON midget cells) during presentation of slow-modulating stimuli in the three stimulus planes. Initially, we observed that the color preferences of OFF midget, ON midget, and blue ON ganglion cells were largely heterogeneous both within and across cell type, even in response to the same color stimulus (Fig. 9). Some clustering of preferred vectors was evident in each stimulus plane: in the (L−M) versus S isoluminant (*xy*) plane (Fig. 9A,B), blue ON cells (*n* = 9) cluster closely to 90° (+S), while both ON (*n* = 19) and OFF midget cells (n = 183) heavily cluster at the two cone-opponent directions near 0° (+L−M) and 180° (−L+M). Notably, a small number of OFF cells show variable preferred vectors depart from the cone-opponent axis toward 270° (−S), with a color signature opposite to blue ON cells (Fig. 9A). While kernel density estimation successfully identified the bimodal distribution of ON and OFF midget cells (OFF: 172° and 2°; ON: 172° and 347°) as well as the unimodal distribution of blue ON cells (101°), the low density of OFF midget cells in the intermediate directions toward −S (225–315°) underlines the overall scarcity of S-OFF cells in our population (Fig. 9B).

**Figure 9.**
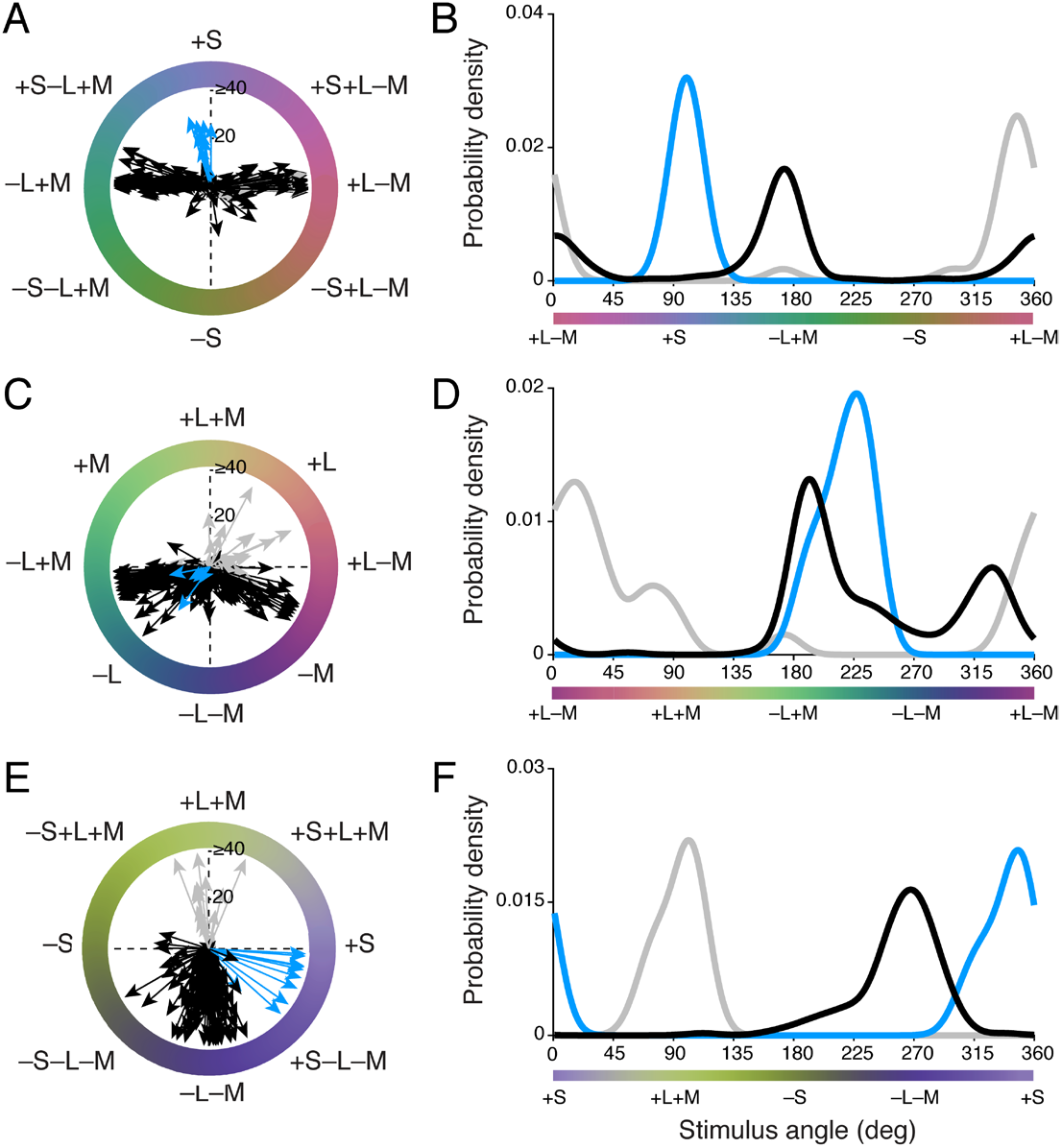
Distributions of color preferences across blue ON, ON midget, and OFF midget ganglion cells. (A–B) Individual preferred vectors (A) and kernel density estimation (B) for cells in the *xy* plane. (C–D) Individual preferred vectors (C) and kernel density estimation (D) for cells in the *yz* plane. (E–F) Individual preferred vectors (E) and kernel density estimation (F) for cells in the *xz* plane. Blue, blue ON cells; gray, ON midget cells; black, OFF midget cells. Vector magnitude is expressed as firing rate (spikes s^−1^) and probability density is normalized for each cell type.

In the (L−M) versus (L+M) (*yz*) plane (Fig. 9C, D), blue ON cells (*n* = 5) cluster at 225°, where the (−L−M) component of these cells is maximally stimulated. ON (*n* = 22) and OFF (*n* = 186) midget cells show a very broad distribution of preferred vectors that span both cone-opponent (0°, 180°) and nonopponent (90°, 270°) directions, illustrating the heterogeneity of opponent responses in midget ganglion cells (e.g., Solomon et al., 2005; Buzás et al., 2006; Martin et al., 2011; Wool et al., 2018). The estimated preferred-vector kernels clearly identified a sharp unimodal distribution of blue ON cells (228°), and a multimodal distribution of ON and OFF midget cells: modes at two L–dominated cone-opponent directions (OFF: 194° and 327°; ON: 174° and 15°), as well as a nonzero density for intermediate, nonopponent L-dominated +L+M (45–90°) and −L−M (225–270°) directions.

In the (L+M) versus S (*xz*) plane (Fig. 9E, F), blue ON cells (*n* = 10) again show a single cluster near 315°, where opponent +S and −L−M mechanisms are maximally stimulated. ON midget cells (*n* = 13) narrowly cluster at 90° (+L+M); by contrast, OFF midget cells (*n* = 143) show a much broader distribution of preferred vectors that span both −L−M and −S mechanisms (Fig. 9E). Compared to the bimodal distributions observed in other stimulus planes, preferred-vector kernels for ON midget, OFF midget, and blue ON cells are unimodal in the *xz* plane, but with varying broadness of their distribution. The preferred-vector kernels for blue ON cells (348°) and ON midget cells (101°) are relatively narrow and normally distributed; by contrast, while the kernel for OFF midget cells shows clear mode at 268°, there is an obvious leftward skew of the distribution, reflecting the substantial number of S-OFF cells that exhibit tuning toward the −S direction (180°) (Fig. 9F).

After computing the preferred vector of each cell to each stimulus (Fig. 9), we computed each cell’s vector projection in three-dimensional color space (Eqs. 7–9 and Methods) and reported this in spherical coordinates: azimuth (*φ*, ranging 0–360° on the *XY* plane) and absolute elevation (*θ*, ranging 0–90°, where 0° lies on the *XY* plane and 90° lies on the *Z* axis). Figure 10 shows the distribution of preferred directions of all ganglion cells in three-dimensional color space. The majority of cells fall in three clusters along the azimuth: blue ON cells fall at *φ* = 90° (+S), while ON and OFF midget ganglion cells fall in columns at *φ* = 0° (+L−M) and *φ* = 180° (−L+M). Moreover, midget cells also show variable tradeoff between being strongly L−M cone-opponent (*θ* = 0°) and nonopponent (*θ* = 90°), consistent with previous studies (e.g., Solomon et al., 2005; Buzás et al., 2006; Field et al., 2010; Martin et al., 2011; Wool et al., 2018). In contrast to the narrow clusters of cells at azimuths of 0°, 90°, and 180°, a sparse population of OFF midget cells are widely scattered between 0° and 180°, indicating a degree of S-OFF input that varies considerably to the receptive fields of these OFF midget cells. These cells also range from strongly chromatic (L−M, *θ* = 0°) to strongly achromatic (L+M, *θ* = 90°); this is consistent with our earlier finding that S-cone inputs are observed across both opponent and nonopponent OFF midget cells.

**Figure 10.**
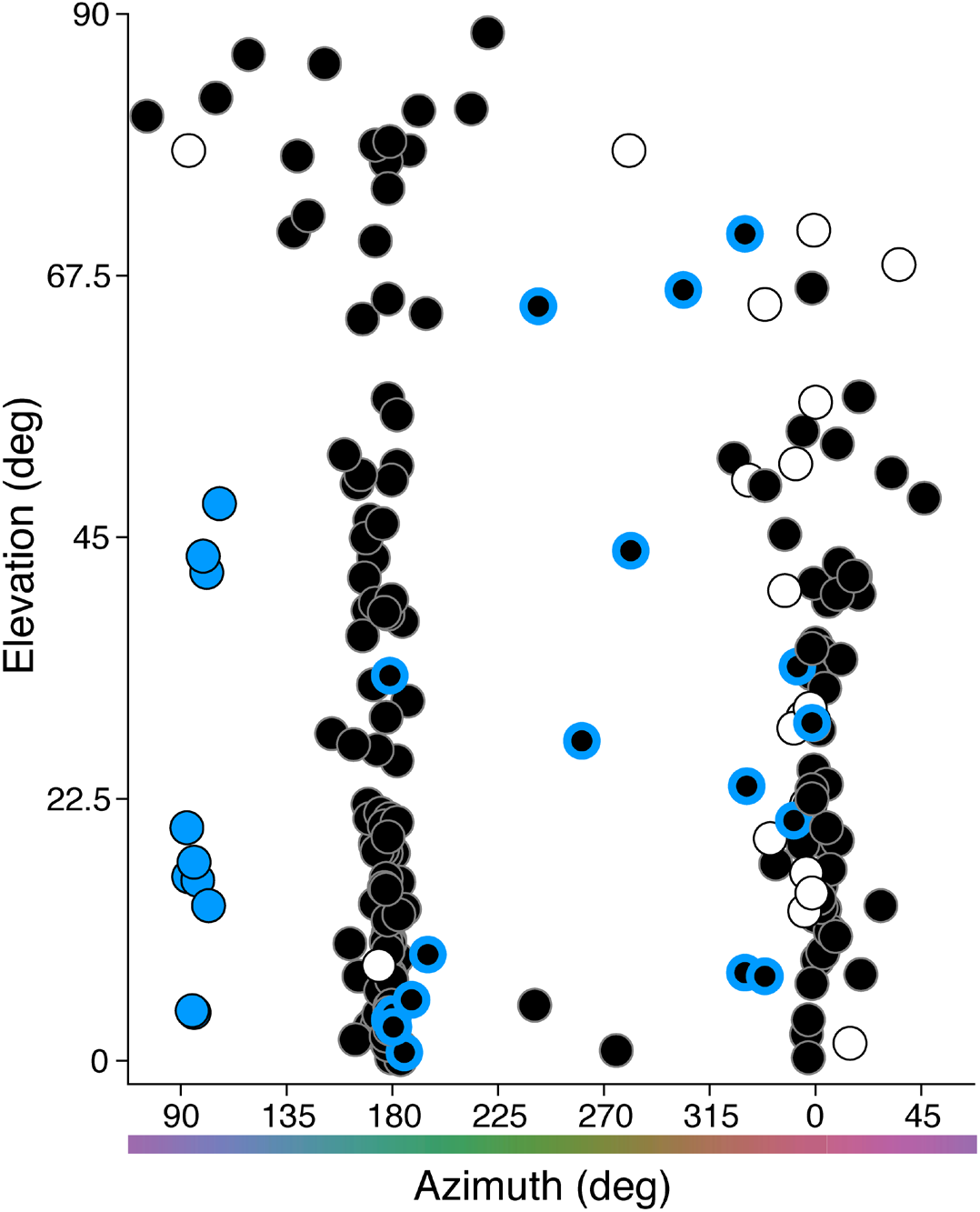
Three-dimensional color signatures of blue ON, ON midget, and OFF midget cells. The preferred azimuth and elevation of each cell was computed from its preferred [*X,Y,Z*] vector in Cartesian coordinates as generated from its preferred vectors across the three individual stimulus planes (see Figure 9). Blue circles, blue ON cells; white circles, ON midget cells; black circles, OFF midget cells. Black/blue circles denote OFF midget cells initially identified as S-cone-sensitive in Figure 1.

## Discussion

By contrast with the S-ON pathway in primate retina, where S cone–selective blue-cone bipolar cells synapse with the small bistratified ganglion cell (Dacey and Lee, 1994; Calkins et al., 1998), a comparable S-OFF pathway does not appear to lie within an anatomically discrete cell class. Instead, our anatomical and physiological results confirm that in the macaque monkey retina, S-OFF signals are propagated by a subpopulation of OFF midget ganglion cells with connectivity to S cones (Calkins, 2001; Klug et al., 2003; Field et al., 2010; Dacey et al., 2014; Tsukamoto and Omi, 2015).

In a previous EM study, Klug et al. (2003) identified five S cones by their connectivity to blue-cone bipolar cells and then showed that each of these cones contacted an OFF midget bipolar cell; two of these cells were completely reconstructed and indistinguishable from OFF midget bipolar cells connected to L/M cones. Thus, the origin of a major S-OFF visual pathway in primate retina appeared to be identified. However, subsequent work in marmoset retina—where S cones were identified by S-opsin expression—showed no such connection to OFF midget bipolar cells (Lee et al., 2005) despite a homologous S-ON small bistratified pathway in this primate species (Ghosh et al., 1997). These results, along with an earlier result suggesting a similar absence of an OFF midget bipolar at presumed S-cone pedicles in the human retina (Kolb et al., 1997) as well as recordings from LGN in both macaque and marmoset identifying a non-midget S-OFF pathway (Szmajda et al., 2006; Tailby et al., 2008a; Tailby et al., 2008b) raised new questions about the retinal substrate for the S-OFF signal in color vision (Lee et al., 2010; Neitz and Neitz, 2011). Later EM reconstructions and physiological data from the retinal periphery (Field et al., 2010; Tsukamoto and Omi, 2015) challenged this apparent absence by again highlighting a sparse S-cone input to the OFF midget circuit.

We took advantage of recent findings that S-cone pedicles uniquely lack the short telodendria that enable L and M cones to interconnect via gap junctions (O’Brien et al., 2012) to unequivocally identify S cone pedicles and to create a larger dataset of S-cone circuits that clearly confirm and extend the result of Klug et al. (2003) (Fig. 11A). Moreover, we found fewer ribbon synapses at both the S-cone pedicle and the associated OFF midget bipolar axon terminal compared to neighboring L/M circuits. The S-cone pathway seems not only distinct in its postsynaptic circuit but also at the quantitative level of ribbon synapse deployment. Given the negative evidence from both human and marmoset, our results suggest significant differences in retinal circuitry across primate groups. It will be critical to use volume EM methods to determine the S-cone connectivity in both marmoset and, most importantly, in human, where connectomics is now being employed to characterize foveal cell types and circuits (Dacey et al., 2017; Packer et al., 2017).

**Figure 11.**
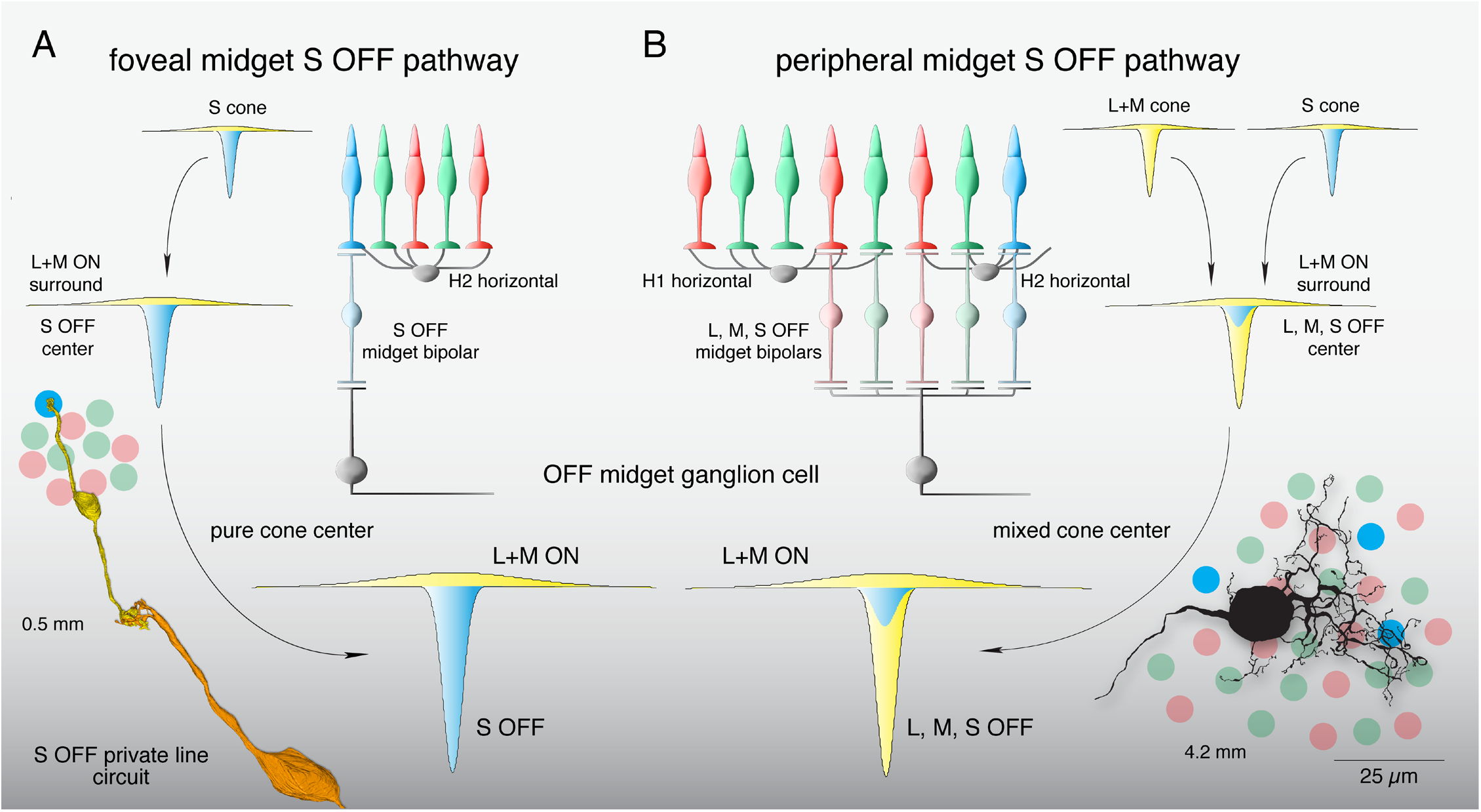
Summary of the OFF midget circuit illustrating a synaptic basis for S-OFF opponent pathways in foveal and near-peripheral retina. (A) An L+M ON surround in the S cone is formed by H2 horizontal cell negative feedback to S cones (Packer et al., 2010). This receptive field structure is transmitted via an S-OFF midget bipolar cell and, subsequently, an OFF midget ganglion cell creating an S-OFF private line circuit (curved arrows). These OFF midget ganglion cells would thus show center-surround organization and S OFF, L+M ON opponency. Inset at lower left shows a schematic foveal cone mosaic of S (blue), L (red), M cone (green) cones in relation to an S-OFF midget circuit reconstruction taken from Figure 3,B–C. (B) In the retinal periphery, L+M ON surrounds are formed in each cone via negative feedback from H1 horizontal cells (to L and M cones) and H2 horizontal cells (to S cones), which are inherited by individual midget bipolar cells and transmitted to midget ganglion cells (curved arrows), which collect input from 3–8 midget bipolar cells in the near retinal periphery (~ 4 mm from the fovea) (Wool et al., 2018). Converging S, L, and M cone inputs create a receptive field in the midget ganglion cell with complex chromatic properties—(S+L) OFF,M ON; (S+M) OFF,L ON; or (S+L+M) OFF—depending on the relative strength of individual cone inputs (Wool et al., 2018). Inset at lower right shows the peripheral cone mosaic, as in panel A, in spatial relation to a near peripheral midget ganglion cell dendritic tree at ~4 mm from the foveal center.

If private-line S-OFF midget circuits were present in the human retina, it would suggest spatial resolution limited by the S-cone sampling array for this pathway. Peak foveal acuity mediated by S-cones is ~10 cpd (Stromeyer et al., 1978; Hess et al., 1989; Metha and Lennie, 2001), exceeding the resolution afforded by Scone spacing. However, acuity drops well below this limit by 2–3° eccentricity (where private-line midget circuitry is present) and this reduction is more consistent with the extensive post-receptoral pooling of the blue ON ganglion cell (Metha and Lennie, 2001), placing it in agreement with other studies of S cone–mediated acuity (Anderson et al., 2002; Beirne et al., 2005). Other measurements suggest that beyond 5°, the S-OFF pathway actually shows lower resolution than its S-ON counterpart (Vassilev et al., 2003; Zlatkova et al., 2008), which is consistent with the rapid post-receptoral pooling of midget ganglion cells that supplants private-line connectivity at greater eccentricities.

In the temporal domain, differences between S-ON and S-OFF signaling do point to their propagation by parallel pathways that sum S cone inputs differently. While some studies observed faster responses to S-OFF stimuli than S-ON stimuli (McKeefry, 2003; Wool et al., 2015), others have reported slower, weaker S-OFF responses both psychophysically (Shinomori and Werner, 2008) and in human electroretinography (Maguire et al., 2018). Such a small and sluggish S-OFF response may be due either to the reduction or absence of this pathway in human retina or to greater variability in the connection strength of the S cones to the OFF midget pathway with increasing eccentricity.

In macaque retina, S-OFF midget cells in the central retina form a private-line pathway but in the periphery the midget cell dendritic tree enlarges and thus draws converging input from L, M, and S cone connecting midget bipolar cells (Jusuf et al., 2006; Tsukamoto and Omi, 2015) (Fig. 11B). Consistent with this picture, we found the midget S-cone signal was variable in strength but invariant in phase: the OFF sign of S-cone sensitivity across OFF-center midget cells is consistent with an S cone contribution to OFF midget ganglion cells via midget bipolar inputs to the excitatory receptive-field center. The spatial extent of S-cone input also supports this interpretation, as the cutoff frequency of spatial tuning curves for S-cone gratings closely matches that of tuning curves to L+M gratings in peripheral OFF midgets. This suggests that in the near periphery where midget dendritic fields receive convergent input from multiple midget bipolar cells, L, M, and S cones all contribute to S-OFF midget ganglion cell receptive field center (Field et al., 2010).

The variable input strength from S cones to OFF midget ganglion cells is consistent with the conclusion that in general, the midget circuit samples nonselectively from the cone mosaic (Calkins and Sterling, 1999; Crook et al., 2011; Wool et al., 2018). Circuit reconstructions in peripheral retina show that S-cone input is variable across neighboring OFF midget ganglion cells, in a manner seemingly dependent on the underlying cone mosaic (Tsukamoto and Omi, 2015). If we assume an S-cone sampling mechanism determined by the sparse S-cone mosaic (Curcio et al., 1991) (and akin to what is observed for L and M cones), we would predict that for midget cells in the far periphery, the likelihood of any S-cone contact would increase, but that the relative strength of S-cone input would decline as overall cone inputs (the great majority L and M) increase. On the other hand, we would predict this tradeoff to be reversed foveally, where a private line OFF midget circuit is present for every S cone, but at a rate in keeping with the low density of S cones. Given that S cones already display an L/M antagonistic surround via negative feedback from horizontal cells (Packer et al., 2010, Fig. 11A), a sparse population of foveal S-OFF center, (L+M) ON surround midget ganglion cells would provide color tuning directly opposing the spatially coextensive S-ON, (L+M) OFF fields of blue ON cells.

We found that the S-OFF connection broadly influences the color-sensitive properties of individual OFF midget cells, giving rise to a population with strongly heterogeneous color preferences that do not lie along the cardinal S vs. L+M and L vs. M chromatic axes (Krauskopf et al., 1982; Derrington et al., 1984). Compared to the consistent S vs. L+M tuning we observed in blue-ON cells, S-OFF midget cells showed L+S OFF, M ON tuning (L and S cones in phase), M+S OFF, L ON tuning (M and S in phase), or (L+M+S) OFF tuning (all cone types in phase). The heterogeneity of phase relationships observed across L, M, and S cone mechanisms in S-OFF midget cells may help to explain why psychophysical studies find that S-cone input can be synergistic with either L-cone input (e.g., Stromeyer et al., 1998; Goddard et al., 2010; Danilova and Mollon, 2012), or M-cone input (e.g., Conway, 2001; Solomon and Lennie, 2005; Horwitz et al., 2007; Lafer-Sousa et al., 2012). These studies measured response amplitudes to modulating ON-OFF gratings assuming symmetry between the poles of various color axes, making it difficult to determine the individual contributions of putative ON and OFF mechanisms to color sensitivity. Our measurements of peak color phase reveal that such poles are rarely symmetric and often driven by multiple cell subtypes (Fig. 5A–C). In any case, our findings suggest that any reported ‘tilt’ off the cardinal L−M axis might be accounted for by the mixed color properties of S-OFF midget cells (Fig. 5A), or their LGN relay cell counterparts (Tailby et al., 2008a).

While the OFF midget circuit may be the dominant pathway for S-OFF signals and set the limit on S-cone spatial resolution, other pathways may also contribute. Melanopsin-expressing intrinsically photosensitive ganglion cells show an S-OFF input (Dacey et al., 2005) and some non-midget diffuse bipolar types also sparsely contact S cones in addition to L and M cones (Calkins and Sterling, 2007; Lee and Grünert, 2007). This suggests additional S-OFF signals involved in color processing may be transmitted to the LGN by ganglion cell types that remain to be characterized (Dacey et al., 2003).

## Acknowledgments

This work was supported by National Institutes of Health grant RR00166 to the Tissue Distribution Program of the National Primate Research Center at the University of Washington, grants EY07556 and EY13312 (to Q.Z.), grant EY06678 (to D.M.D.), and grant EY01730 (to the Vision Research Core at the University of Washington). We thank Julian Vrieslander for programming assistance and Beth Peterson, Ed Parker, Dale Cunningham, Yeon Jin Kim, and Ursula Bertram for their contributions. Territorial acknowledgments: This study was undertaken at institutions occupying unceded Indigenous land of the Coast Salish people (specifically the Duwamish Tribe) (University of Washington) and the Lenape people (SUNY College of Optometry). The authors declare no competing financial interests.

